# Restoration of Kv7 channel mediated inhibition reduces cued-reinstatement of cocaine seeking

**DOI:** 10.1101/272971

**Authors:** Jeffrey Parrilla-Carrero, William C. Buchta, Priyodarshan Goswamee, Oliver Culver, Greer McKendrick, Benjamin Harlan, Aubin Moutal, Rachel Penrod, Rajesh Khanna, Peter Kalivas, Arthur C. Riegel

**Author notes:** Equal contribution (Jeffrey Parrilla-Carrero, William C. Buchta, Oliver Culver, Greer McKendrick). Correspondence, Department of Neurosciences, Medical University of South Carolina (MUSC), 410C Basic Sciences Building. Author contributions: J.P.C, W.B., P.G, O.C., B.H., A.M., R.K., P.K., and A.C.R. designed research; J.P.C, W.B., P.G, O.C., B.H., A.M., R.P., and A.C.R. performed research; J.P.C, W.B., P.G, O.C., G.M., B.H., A.M., R.P., R.K., P.K., and A.C.R. analyzed data; J.P.C, W.B., and A.C.R. wrote the paper.

## Abstract

Cocaine addicts display increased sensitivity to drug-associated cues, due in part to pathological changes in the prelimbic cortex (PL-PFC). The cellular mechanisms underlying cue-induced reinstatement of cocaine seeking remain unknown. Reinforcement learning for addictive drugs may produce persistent maladaptations in intrinsic excitability within sparse subsets of PFC pyramidal neurons. Using a male rat model of relapse to cocaine-seeking, we sampled over 600 neurons to examine spike frequency adaptation (SFA) and after-hyperpolarizations (AHPs), two systems that attenuate low frequency inputs to regulate neuronal synchronization. We observed that training to self-administer cocaine or nondrug (sucrose) reinforcers decreased SFA and AHPs in a sub-population of PL-PFC neurons, but only with cocaine did the resulting hyper-excitability persist through extinction training and increase during reinstatement. In neurons with intact SFA, dopamine enhanced excitability by inhibiting Kv7 potassium channels that mediate SFA. However, dopamine effects were occluded in neurons from cocaine-experienced rats, where SFA and AHPs were reduced. Pharmacological stabilization of Kv7 channels with retigabine restored SFA and Kv7 channel function in neuroadapted cells. When microinjected bilaterally into the PL-PFC 10 minutes prior to reinstatement testing, retigabine reduced cue-induced reinstatement of cocaine seeking. Lastly, using cFos-GFP transgenic rats, we found that the loss of SFA correlated with the expression of cFos-GFP following both extinction and re-exposure to drug-associated cues. Taken together, these data suggest that cocaine self-administration desensitizes inhibitory Kv7 channels in a subpopulation of PL-PFC neurons. This sub-population of neurons may represent a persistent neural ensemble responsible for driving drug seeking in response to cues.

**Significance Statement:** Long after the cessation of drug use, cues associated with cocaine still elicit drug-seeking behavior, in part by activation of the prelimbic cortex (PL-PFC). The underlying cellular mechanisms governing these activated neurons remain unclear. Using a rat model of relapse to cocaine seeking, we identified a population of PL-PFC neurons that become hyperexcitable following chronic cocaine self-administration. These neurons show persistent loss of spike frequency adaptation, reduced after-hyperpolarizations, decreased sensitivity to dopamine, and reduced Kv7 channel mediated inhibition. Stabilization of Kv7 channel function with retigabine normalized neuronal excitability, restored Kv7 channel currents, and reduced drug-seeking behavior when administered into the PL-PFC prior to reinstatement. These data highlight a persistent adaptation in a subset of PL-PFC neurons that may contribute to relapse vulnerability.

## Introduction

The prefrontal prelimbic cortex (PL-PFC) plays an important role in relapse to cocaine seeking. This brain region filters input from sensory and limbic areas and is critical for initiating appetitive behaviors via its projections to the basal ganglia (Graybiel, 2008). Chronic exposure to drugs of abuse causes pathological changes in the PL-PFC and other learning and memory-related brain structures that contribute to relapse (Hyman et al., 2006; Williams and Adinoff, 2007). Specifically, after extinction from chronic cocaine self-administration, activated glutamatergic efferents from the PL-PFC innervating the nucleus accumbens drive cue-induced reinstatement of drug-seeking (Kalivas and McFarland, 2003; See, 2009; Stefanik et al., 2016).

Behaviorally relevant cues activate only a small portion of neurons within a brain region (Cruz et al., 2013a; 2015). Across many brain regions, these sparse ensembles of activated neurons contribute to the processing of sensory cues and are thought to store the memory trace for such learned associations (Matsumoto and Hikosaka, 2009; Euston et al., 2012; Pinto and Dan, 2015). These neuronal ensembles of activated neurons can be identified using immediate early gene indicators like cFos, whose expression increases following strong neural activation (Morgan et al., 1987; Morgan and Curran, 1988; Herrera and Robertson, 1996). For example, following cue-induced reinstatement, PL-PFC neurons that project to the accumbens core show robust cFos activation, while cells that project to the accumbens shell do not. This PL-PFC to accumbens core pathway is recruited in a dopamine-dependent manner to drive drug seeking in response to cues (McGlinchey et al., 2016). Dopamine signaling can induce cFos expression (Zhang et al., 2006) and promote the formation and maintenance of activity within cellular neural ensembles (O’Donnell, 2003; Puig and Miller, 2012a; Miyawaki et al., 2014). However, the cellular adaptations that occur within these activated ensembles in the PL-PFC remain unclear.

At the cellular level, spike frequency adaptation (SFA) and after-hyperpolarizations (AHPs) are two principle forces regulating neuronal synchronization (Crook et al., 1998; Prescott and Sejnowski, 2008; Prescott et al., 2008). SFA decreases responses to non-preferred stimuli (Benda et al., 2005; Simon Peter Peron, 2009). However, SFA is typically dynamic and reduced by many neuromodulators. For example, we recently showed that dopamine release from ventral tegmental area (VTA) terminals in the PL-PFC can reduce SFA (Buchta et al., 2017). In other brain regions, reduced SFA in ensembles of cells can enhance memory storage and behavioral responses to cues encoded by the ensemble (Yiu et al., 2014). Across multiple brain regions and behavioral paradigms, reduced SFA is associated with learning (Moyer et al., 1996; McKay et al., 2009; Sehgal et al., 2014). Relative to non-drug reinforcers such as sucrose, drugs of abuse produce robust, long-lasting neural changes that result in stronger, more persistent behavioral responding for drugs (Ciccocioppo et al., 2004; Tunstall and Kearns, 2016). Thus, reinforcement learning for drugs may produce persistent adaptations in SFA that contribute to enduring cue-reward associations and the subsequent ability of cues to elicit drug-seeking behavior. However, whether SFA is disrupted in the PL-PFC following chronic cocaine self-administration and how this contributes to relapse to drug seeking have not been previously examined.

Here, we used *ex vivo* electrophysiology to examine SFA following cocaine self-administration training, extinction, and reinstatement. We assessed whether previous cocaine self-administration experience modified the responses of activated neurons to dopamine. We also examined the function of the Kv7 potassium channels contributing to SFA and assessed whether dysfunction of these channels contributed to drug-seeking behavior in response to cues. Lastly, we used cFos-GFP transgenic rats to identify and record from cFos-GFP^+^ neurons and examine whether SFA changes were specific to GFP^+^ neurons. Our results identify a subset of cFos-GFP^+^ PL-PFC neurons that show a persistent loss of SFA and Kv7 channel inhibition following chronic cocaine self-administration. We further show that pharmacological stabilization of Kv7 channels with retigabine restores SFA, Kv7 channel inhibition, and reduces cue-induced reinstatement. These cells may represent an enduring neural ensemble that maintains drug-cue associations and drives drug seeking in response to cues.

## Materials and Methods

### Animals

We used adult (>P65) male Sprague Dawley (Charles River Laboratories, Indianapolis, IN) and transgenic cFos-GFP Long Evans (bred in-house; breeding pair provided by Dr. Bruce Hope) rats, each weighing between 250 and 300 g at the beginning of experiments. Rats were individually housed in a temperature and humidity-controlled environment under a reversed 12-h light/dark cycle (lights on at 6:00 PM). Rats were acclimated for a minimum of 1 week prior to surgeries and had access to food and water *ad libitum*. All procedures, sacrifices, and behavioral experiments were performed during the dark cycle, and in accordance with guidelines obtained from the National Institutes of Health (NIH) for the care of laboratory animals and by the Medical University of South Carolina (MUSC) Institutional Animal Care and Use Committee (IACUC).

### Surgery

Surgical procedures were performed on rats to implant catheters for receiving intravenous cocaine or saline infusions, or intra-cranial canulae for injection of pharmacological compounds directly to the prelimbic cortex. Rats were anaesthetized using a ketamine HCl / xylazine mixture (0.57 / 0.87 mg/kg, respectively, IP) followed by ketorolac (2.0 mg/kg, IP) and Cefazolin (40 mg, IP or 10 mg / 0.1 ml, IV). Subsequently, intra-jugular catheters were implanted as described elsewhere (Spencer et al., 2017). Catheter patency was maintained by daily flushing with cefazolin (1 g / 5 ml during surgery and 1 g / 10 ml daily for 1 week) and then with heparin for the remainder of the behavioral paradigms. Some rats received bilateral intracranial microinjections of either retigabine or vehicle (dimethyl-sulfoxide, DMSO) via canulae (double 28ga barrel; 1.2-1.5 mm C-C; Plastics One) that were stereotactically implanted into the brain and reached 1 mm above PL-PFC using the following coordinates relative to Bregma (in mm; AP: + 3.0-3.5; DV: -3.0; ML: ± 0.75).

### Behavioral procedures

The rationale underlying the self-administration, extinction and cued-reinstatement testing is described in detail elsewhere (Steketee and Kalivas, 2011; Bossert et al., 2013). The general timeline outlining the different behavioral phases is shown in Figure 1*A*. For experiments 1, 2, 4, and 5, rats were trained to self-administer cocaine (coca-SA group) on a fixed ratio (FR) 1 schedule of reinforcement (2h / session). Intravenous injection of cocaine (∼0.2 mg/kg/infusion) was delivered contingent to pressing one (active) of two levers in response to a compound light (5 s) and tone (4 KHz, 78 dB, 5 s) cue that was followed by a 20 s time-out period. Each training session lasted 2 h, after which animals were returned to their home cages. A parallel set of time-matched control rats were “yoked” to the cocaine self-administration group and received either saline (sal-Y group) or non-contingent infusions of cocaine (coca-Y group), every time the paired cocaine self-administration rat pressed the active lever during the self-administration phase (Figure 1*A)*.

**Figure 1.**
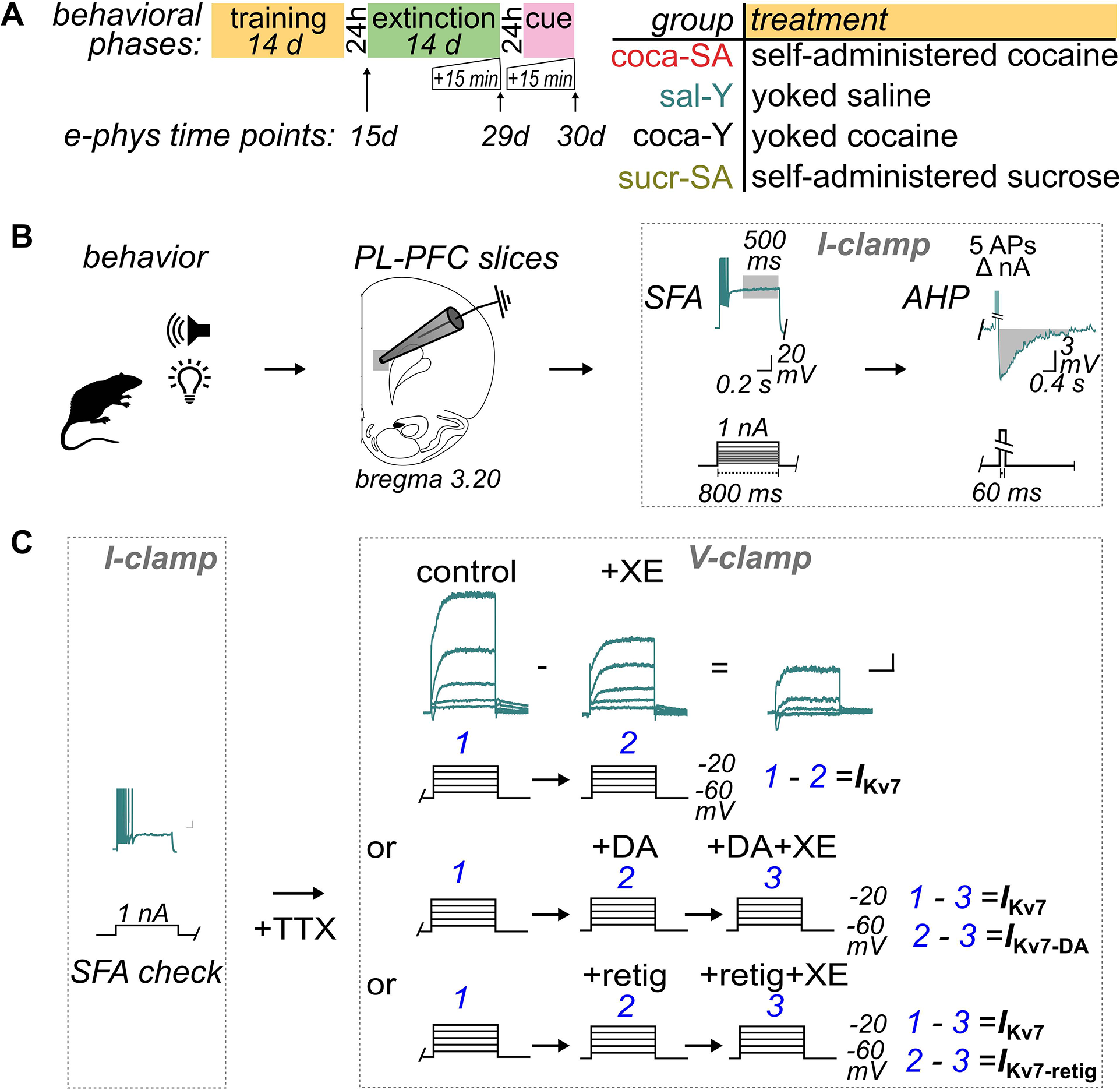
Experimental outline for behavioral training and physiological measurements. ***A*** Time line shows the different behavioral phases, time points of electrophysiological measurements (e-phys), and the color-coded treatment groups. ***B*** After behavioral end points, brain slices containing prelimbic (PL) prefrontal cortex (PFC) were made for current clamp measurements of SFA (measured during the final 500 ms of the step) and AHPs generated in response to 5 action potentials (APs) (expressed as the area under the curve). ***C*** In some experiments, after measuring firing to check for SFA with a 1 nA step, the *I*_Kv7_ was determined in voltage clamp. Currents were evoked with voltage steps (-60 to -20 mV; 800 ms; in 200 nM TTX). Currents generated after 8 min incubation of the Kv7 channel antagonist, XE-991 (20 μM, 8 min) were subtracted from control (pre-drug) currents. To determine the effect of dopamine (DA, 10 μM) or the Kv7 channel stabilizer retigabine (retig, 20μM) on *I*_Kv7_, the currents generated after bath application of DA + XE-991 (20 μM, 8 min) or retigabine + XE-991 (20 μM, 8 min) were subtracted from currents after DA or retigabine alone, respectively.

**Figure 2.**
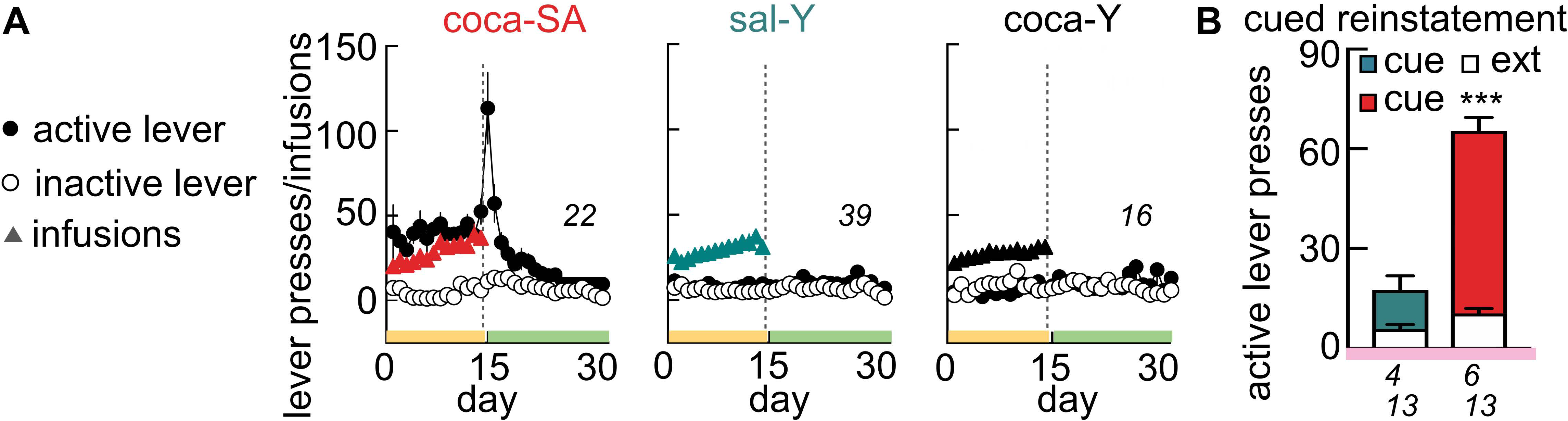
Behavioral responding in cocaine self-administration, yoked-cocaine and yoked-saline rats. ***A*** Rats were presented with levers that resulted in activation of a light and tone cue paired with infusion of cocaine (coca-SA, *left*), passive infusion of saline (saline, sal-Y, *middle*), or passive infusion of cocaine (coca-Y, *right*). The vertical dashed line on time courses indicates the switch from self-administration to extinction. In extinction sessions, lever pressing produced no infusions or cue presentations. Behavioral responses for voluntary cocaine self-administration extinguished in 14 days. ***B*** Re-exposure to drug-conditioned cues (without cocaine) significantly increased lever pressing in rats with a history of coca-SA. Numbers in *italics* represent the total number of animals. For these and all other figures, error bars indicate mean ± SEM. *p≤0.0001 compared with extinction using a Sidak test for multiple comparisons.

#### Experiment 1 (Figure 3)

Following maintenance of stable self-administration behavior for 14 consecutive days (>10 infusions / session), the sal-Y, coca-Y and coca-SA rats were sacrificed 24 hrs after the last training session to harvest brain tissue for electrophysiological recordings.

**Figure 3.**
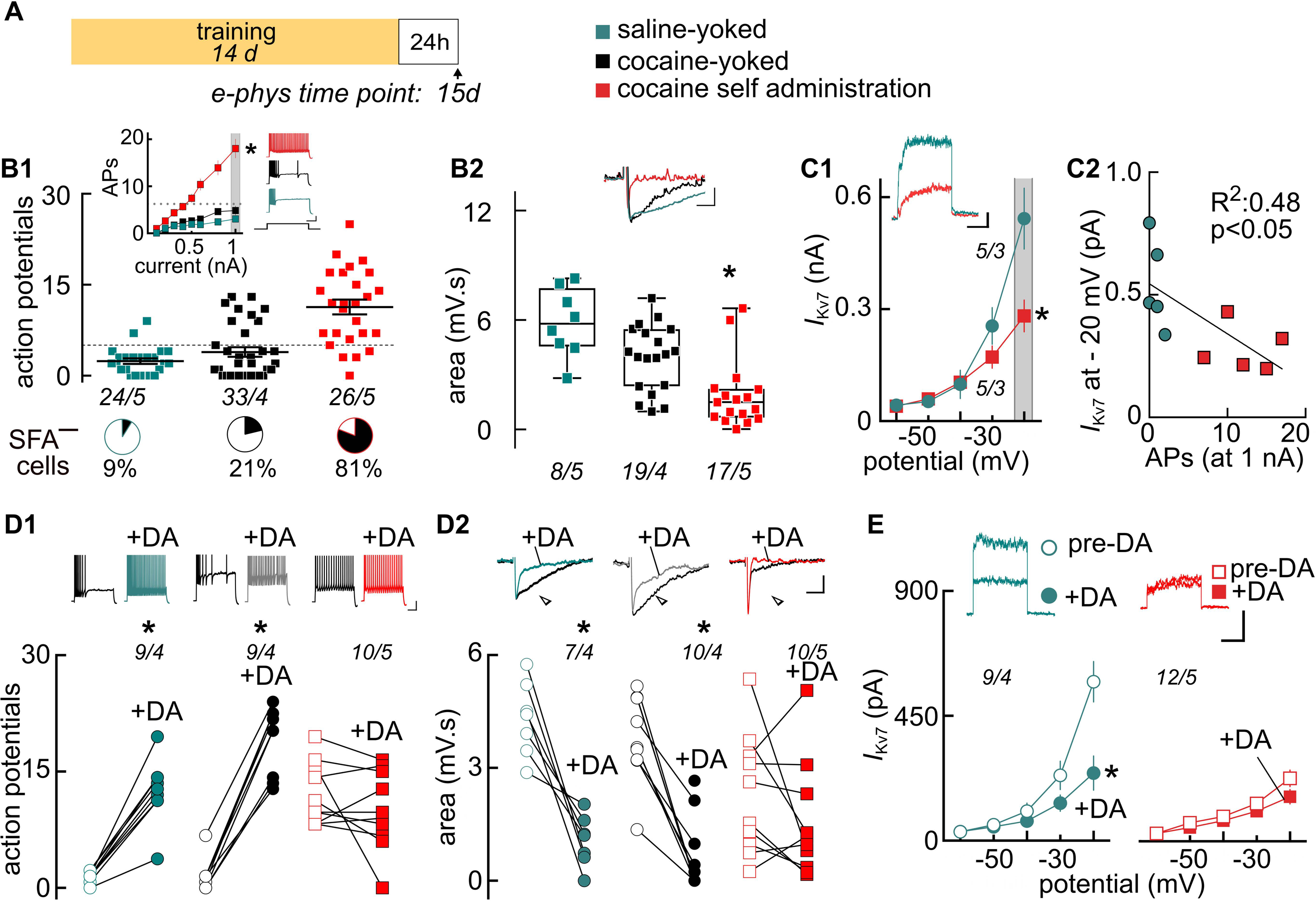
Experiment 1: Training to self-administer cocaine (but not passive exposure to cocaine) suppressed intrinsic inhibition and inhibitory Kv7 channel currents. ***A*** Timeline showing behavioral training ended 24 hr prior to sacrifice for electrophysiological recordings. ***B1*** Scatter plots show firing responses of individual cells at 1nA. Matching I-O curves (*inset*) show increased mean firing responses for the coca-SA relative to coca-Y or sal-Y treatment groups. Firing exceeded the upper limit for SFA (operationally defined as ≤ 5 spikes at 1 nA, *dashed grey line*) only in the coca-SA group. Black segments in the pie charts indicate % of sampled cells that were negative for SFA (SFA ^―^). ***B2*** Scatter plots show decreased average AHPs in the coca-SA group only. ***C1*** I-O curves show reduced *I*_Kv7_ currents in the coca-SA group. Prior to *I*_Kv7_ measurements, cells in the coca-SA and sal-Y groups were confirmed to be SFA ^―^ and SFA^+^, respectively. ***C2*** Reductions in *I*_Kv7_ current amplitudes correlated with action potential (AP) firing at the 1 nA step. ***D1,2*** Dot plot graphs show dopamine increased firing (D1) and decreased AHP (D2) in coca-SA/ SFA^+^ and sal-Y/ SFA^+^ cells. Dopamine produced no measurable change in coca-SA/SFA ^―^ cells. ***E*** Dopamine reduced *I*_Kv7_ in the sal-Y/SFA^+^ but not in the coca-SA/SFA ^―^ cells. All *insets* show sample traces collected at 1 nA step for firing (I-clamp) or at -20mV for Kv7 (V-clamp). *Italicized* numbers represent the number of cells / animals. Scale bars: B1: 20 mV, 0.2 s; B2: 3 mV, 0.4 s; C1: 0.1 nA, 0.4s, D1: 20 mV, 0.2 s, D2: 3 mV, 0.5 s; E: 0.1 nA, 0.4 s. ^*^p≤0.01 compared with sal-Y using a Sidak test for multiple comparisons (Figures 3*B,C,E)* or paired *t*-test against pre-DA baselines (Figure 3*D*).

#### Experiment 2 (Figure 4)

24 hrs after the last cocaine self-administration (or yoked) training session, animals entered into extinction training to extinguish lever-pressing behavior. During the extinction period (14-19 days) pressing of the active lever yielded neither cocaine nor the light/tone cue. Once the self-administration behavior was stably extinguished in coca-SA rats (<25 active lever presses for 3 consecutive days), sal-Y and coca-SA rats were sacrificed 15 min after completion of their last extinction session to harvest brain tissue for electrophysiological recordings.

**Figure 4.**
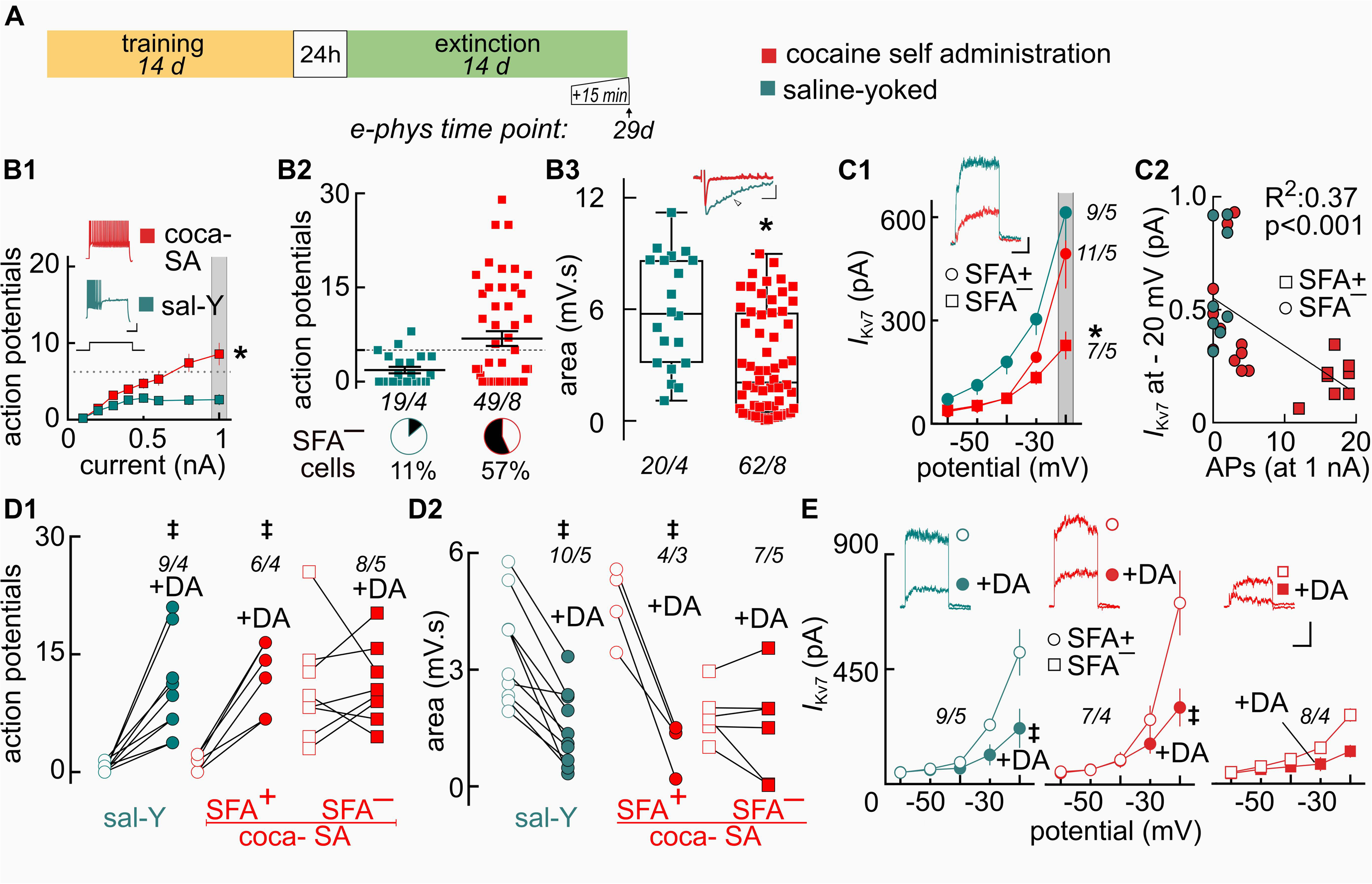
Extinguishing the behavioral responding for cocaine does not restore intrinsic inhibition or Kv7 channel currents. ***A*** Time line showing cocaine-SA (14d) and extinction (14d) prior to sacrifice for electrophysiological recordings at 15 min after the final the final extinction session. ***B*** Average firing (B1, I-O curves) and individual firing (B2, scatter plots at 1 nA) was increased in the coca-SA group, while AHP (B3, scatter plots) was decreased relative to sal-Y controls. Black segments in the pie charts indicate % of sampled cells that were negative for SFA (SFA ^―^). ***C1*** I-O curves show reduced *I*_Kv7_ currents in the coca-SA/SFA ^―^ cells, but not the sal-Y/SFA^+^ or coca-SA/SFA^+^ cells. ***C2*** Reductions in *I*_Kv7_ current amplitudes correlated with action potential (AP) firing at the 1 nA step. ***D*** Dot plot graphs show dopamine increased firing (D1) and decreased AHPs (D2) in SFA+ cells from both sal-Y and coca-SA groups. Dopamine did not measurably alter coca-SA/SFA ^―^ cells. ***E*** Dopamine reduced *I*_Kv7_ in SFA+ cells of sal-Y and coca-SA rats, but not in SFA ^―^ cells. All *insets* showing sample traces were collected at 1 nA step for firing (I-clamp) or at -20mV for Kv7 (V-clamp). *Italicized* numbers represent the number of cells / animals. Scale bars: B1: 20 mV, 0.2 s; B3: 3 mV, 0.4 s; C1: 0.1 nA, 0.4s, D1: 20 mV, 0.2 s, D2: 3 mV, 0.5 s; E: 0.1 nA, 0.4 s. *p ≤ 0.001 compared with sal-Y using a Sidak test for multiple comparisons (Figures 4*B,C,E)* or paired *t*-test against pre-DA baselines (‡, Figure 4*D*).

#### Experiment 3 (Figure 5)

After the last cocaine self-administration session, 6 coca-SA rats were returned and remained in their homecage for the remainder of a 14d withdrawal period (Figure 5*A*). Rats were sacrificed on the last day of withdrawal for electrophysiological recordings. Another group of 5 unhandled and behaviorally naïve rats were also sacrificed for brain slices (Figure 5*B*). A third group of 12 rats was trained to self-administration sucrose (sucrose-SA; Figure 5*C*) using a procedure similar to previously published studies (McGlinchey et al., 2016), where rats were handled daily for at least 1 week before sucrose self-administration training. Rats were trained to self-administer sucrose on an FR1 schedule of reinforcement (2h / session) in operant chambers within sound-attenuating boxes, controlled by Med-PC IV software (Med Associates). Sucrose pellets (45 mg, Test Diet) were delivered contingent to pressing one (active) of two levers in response to a compound light (5 s) and tone cue (4 KHz, 78 dB, 5 s) followed by a 20 s time-out period, where additional presses had no programmed consequences. Presses on the inactive lever were recorded but had no programmed consequences. After 10-14 d of criterion sucrose-SA performance (≥30 pellets / session), one group of 6 rats were sacrificed 24 hrs after the last training session to harvest brain tissue for electrophysiological recordings. The remaining 6 rats underwent extinction training, where responding on either lever had no consequence. Rats received extinction training for a minimum of 10 d and until they met the criteria of <25 active lever presses for 3 or more consecutive days. Rats were sacrificed 15 min after the last extinction session to harvest brain tissue for electrophysiological recordings.

**Figure 5.**
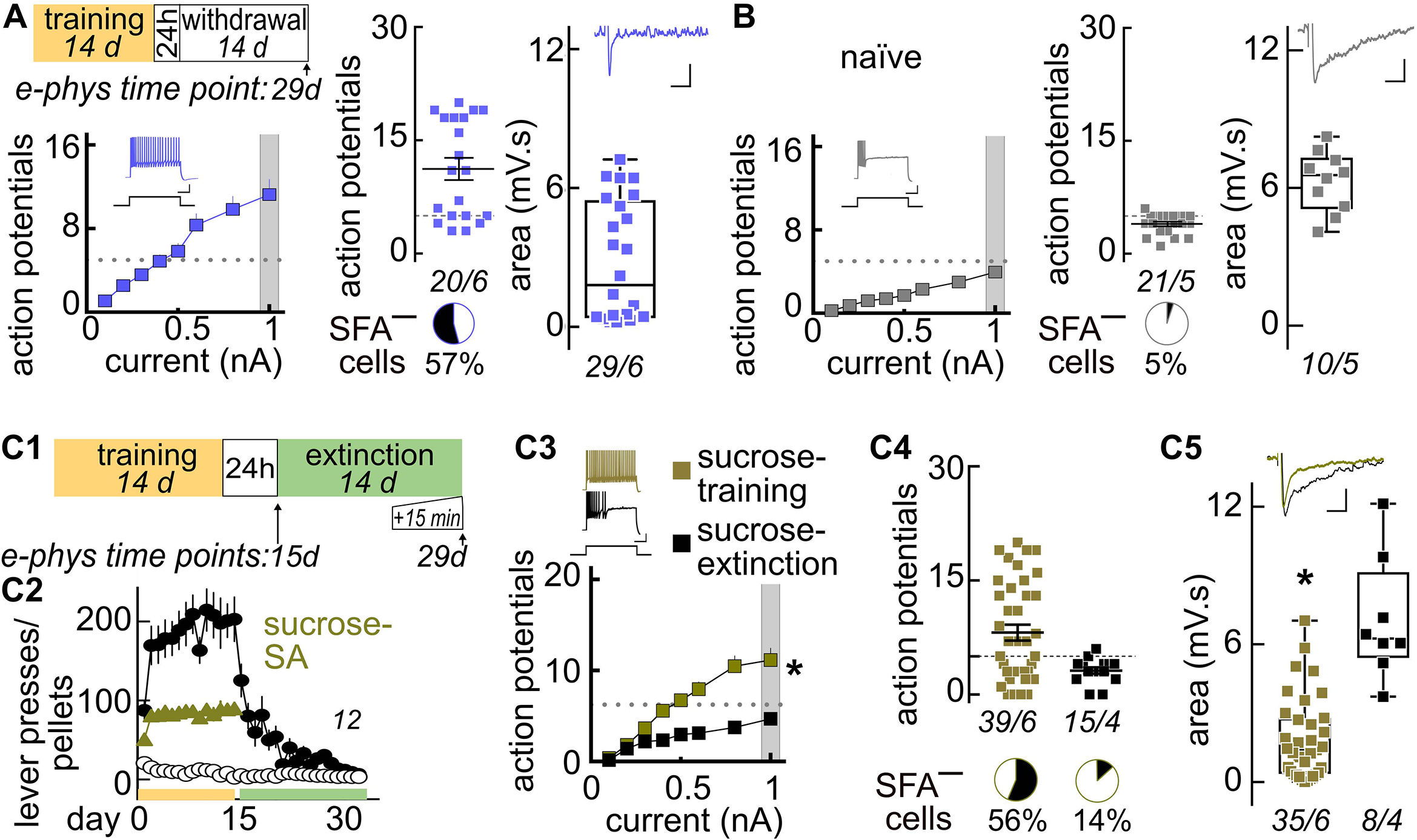
Differences in intrinsic inhibition following withdrawal from cocaine-SA and behavioral extinction from sucrose-SA. ***A*** *Top* Timeline showing cocaine-SA with 14d of home cage withdrawal, ending immediately prior to the preparation of brain slices for electrophysiological recordings. Cells from rats withdrawn from cocaine-SA (without extinction training) showed elevated average firing beyond the maximal range for SFA (I-O curve with *inset* sample trace collected at 1 nA), heterogeneous individual firing responses (scatter plots at 1 nA), and reduced AHPs (*inset* sample trace and scatter plots). ***B*** Responses from unhandled, experimentally naÏve rats showed robust SFA across the I-O curve and homogenous individual firing (scatter plots at 1 nA) and AHP (scatter plots) responses. ***C1*** Timeline showing 14d of sucrose-SA followed by extinction (14d). Preparation of brain slices occurred at either 24 hr after the last SA session (d15) or 15 min after the final extinction session (d29). ***C2*** Rats were presented with levers that resulted in activation of a light and tone cue paired with delivery of a sucrose pellet. The vertical dashed line on time courses indicates the switch from self-administration to extinction. In extinction sessions, lever pressing had no consequence. Behavioral responses for sucrose extinguished in 14 days. ***C3-5*** Comparison of responses showing the increased excitability associated with sucrose-SA resolves with extinction, including average firing (C3, I-O curves), individual firing responses (C4, scatter plot, at 1 nA) and AHPs (C5, scatter plot). The black segments in pie charts below indicate % of sampled cells that were negative for SFA (SFA ^―^). *Italicized* numbers indicate the number of cells / animal. Scale bars: Figure 5*A, B, C*: firing traces, 20 mV, 0.2 s; AHP traces, 3 mV, 0.4 s. *p≤0.001 compared with sal-Y using a Sidak test for multiple comparisons or unpaired *t*-test.

#### Experiment 4 (Figure 6)

Once cocaine self-administration was completed and behavioral responding was stably extinguished (<25 active lever presses for 3 consecutive days), sal-Y and coca-SA rats returned to the home cage for 24 hrs. The next day, they were tested for cue-induced reinstatement of lever pressing (Epstein et al., 2006). During the 2 hr test, rats were re-exposed to the cocaine-paired cues (but, not cocaine) with a 20 s timeout period after each press of the active lever. Rats were sacrificed immediately at the end of the 2 hr test for electrophysiological recordings. In a separate set of coca-SA rats (n= 27 rats), retigabine (0.0015 /0.015 / 0.15 nmol / 0.5 μl prepared in saline with 0.1% DMSO) or vehicle (0.1% DMSO in 0.5ul saline) was microinjected into the PL-PFC 10 min prior to 2 hr cued-induced reinstatement sessions. A randomized counterbalanced design with 2-3 cue reinstatement tests was used, with at least 2 days of extinction (≤25 active lever presses) between tests (Smith et al., 2014). Following the completion of reinstatement testing, locomotor testing was conducted in novel open-field chambers (AccuScan Instruments, Inc.). PL-PFC infusions (identical as described above) of either retigabine or vehicle were made 10 mins prior to placement in the open-field chambers. Total distance traveled in the following 2 hrs was quantified using the Versamax software. Data were included for analysis only if the histology identifying the center of injection sites confirmed PL-PFC cannulas placements, which ranged from ∼3.2-4.2 mm AP.

**Figure 6.**
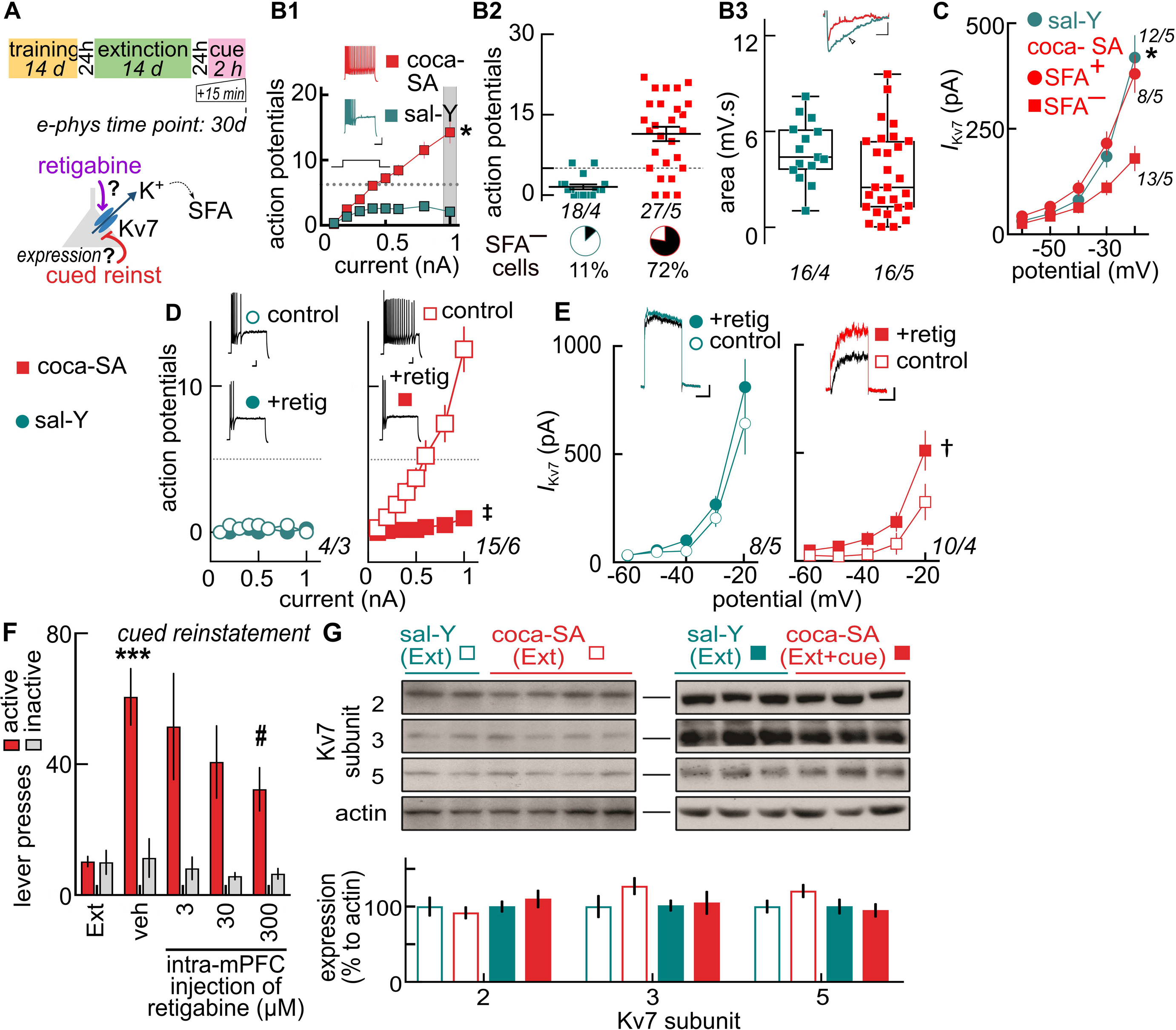
Cues that reinstate cocaine seeking rekindle the suppression of intrinsic inhibition: Restoration of inhibition by stabilization of Kv7 channels reduces cocaine seeking. ***A*** Timeline showing the preparation of brain slices, occurring 15 min after the 2 hr cued-reinstatement test. Cartoon shows proposed retigabine action on *I*_Kv7_ in SFA ^―^ cells during reinstatement. ***B*** Comparison of responses showing cued-reinstatement reduces intrinsic inhibition in coca-SA treatments relative to sal-Y controls. Shown are average firing (B1, I-O curves), individual firing responses (B2, scatter plot, at 1 nA) and AHPs (B3, scatter plot). Black segments in pie charts below indicate % of sampled cells that were negative for SFA (SFA ^―^). ***C*** Relative to SFA+ (coca-SA or sal-Y) cells, *I*_Kv7_ currents are reduced in SFA ^―^ (coca-SA) cells. ***D,E*** Summary graphs showing retigabine (retig, 20μM) reduced firing (D) and increased *I*_Kv7_ (E) only in SFA ^―^ (coca-SA) cells, and not in SFA+ (sal-Y and coca-SA) cells. ***F*** Bilateral microinjection of retigabine into PL-PFC 10 min prior to reinstatement testing resulted in a dose-dependent reduction of cue-induced reinstatement. ***G*** Western blots of PL-PFC punch lysates showing Kv7 subunit expression (7.2, 7.3, 7.5) as a percentage of actin control did not differ between sal-Y and coca-SA, either after extinction or after cued-reinstatement. Scale bars: B1 and D: 20 mV, 0.2 s; B3: 3 mV, 0.4 s; C1 and E: 0.1 nA, 0.4 s. All *insets* showing sample traces were collected at 1 nA step for firing (I-clamp) or at -20mV for Kv7 (V-clamp). *Italicized* numbers indicate the cells / animal. Sidak test for multiple comparisons: p ≤ 0.0001 compared with sal-Y (*, Figure *B1, C*) or pre-retigabine (‡, Figure *D*); p ≤0.01 compared with pre-retigabine (‡, Figure *E*). (F) ***p≤0.0001 vs ext, #p=0.0299 veh vs retig (300uM).

#### Experiment 5 (Figure 7)

Transgenic cFos-GFP rats were trained to self-administer cocaine (coca-SA group) using the same procedure as described above. A parallel set of time-matched control rats were “yoked” to the coca-SA group, but received saline (sal-Y group) every time the paired coca-SA rat pressed the active lever during the self-administration phase. Following the same procedure as in experiment 1, 4 sal-Y and 5 coca-SA rats were sacrificed immediately after the last 90 min extinction session to harvest brain tissue for electrophysiological recordings. The remaining 4 sal-Y and 5 coca-SA rats were extinguished as described in Experiment 4 and then tested for cued-reinstatement of lever-pressing as in Experiment 4, except that the session lasted 90 min. Rats were sacrificed immediately at the end of the 90 min cue test for electrophysiological recordings.

**Figure 7.**
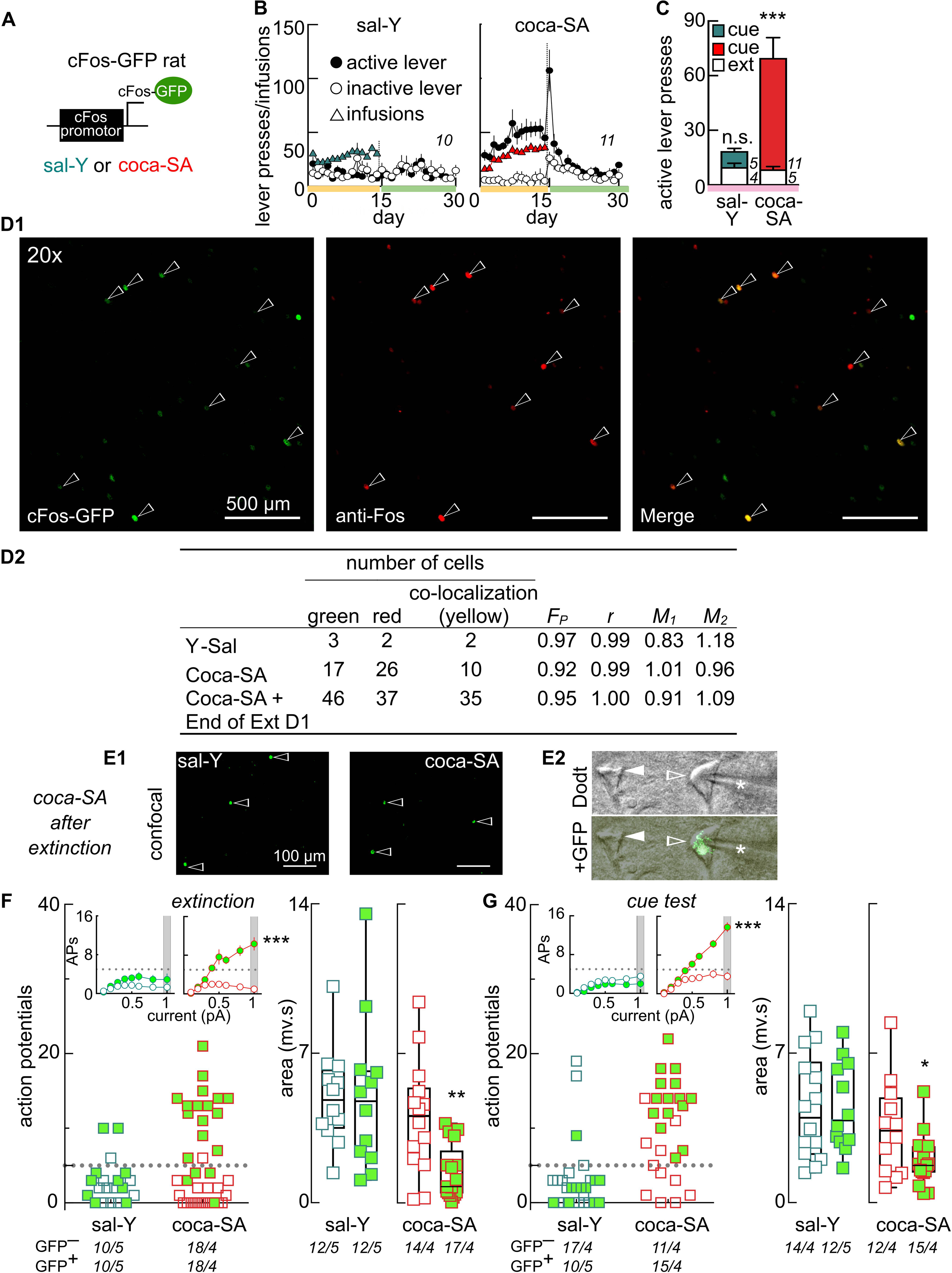
Characterization of intrinsic inhibition in cFos-GFP^+^ and cFos-GFP ^―^ cells in the PL-PFC after extinction and cued-reinstatement of cocaine seeking. ***A*** Schematic of the transgene containing a c-fos promoter that induces green fluorescent protein (GFP) in strongly activated Fos-expressing neurons, which can be sampled in electrophysiological slice preparations. ***B*** Lever pressing behavior for Long Evans rats providing the PL-PFC electrophysiology recordings and immunohistochemistry in Figure 7. Rats were presented with levers that resulted in activation of a light and tone cue paired with infusion of cocaine (coca-SA, *right*) or yoked passive infusion of saline (sal-Y, *left*). The vertical dashed line on the time courses indicates the switch from self-administration (SA) to extinction (EXT). In EXT sessions, lever pressing had no consequence. ***C*** Re-exposure to cues (without saline or rewards) significantly increased lever pressing in rats with a history of coca-SA (Sidak post hoc, ***p≤0.0001, cue vs extinction). ***D*** Confocal photomicrographs (D1) showing colabeling (white arrows) of GFP (green) and Fos (red) using immunofluorescence histochemistry. Micrographs are from a c-Fos-GFP rat that underwent coca-SA and a single EXT training session prior to perfusion (d15). (D2) Table summarizing GFP/Fos co-colabeling in transgenic rats with different treatments: Rat 1, sal-Y+14d EXT(sacrifice d28); Rat 2-coca-SA(sacrifice d14); Rat 3 coca-SA+1d EXT(sacrifice d15). A high degree of colocalization between fluorophores was evident from the Pearson’s Correlation Coefficient (*F*_*P*_; summarizing the ratio of cells that colocalize) and Manders’ Overlap Coefficient (*r*; showing the overlap between channels). ***E1*** Confocal photomicrographs show similarly low levels of GFP-IR (open arrowheads) in the PFC of a sal-Y (*left*) and coca-SA (*right*) rat immediately after the final EXT session (d28). ***E2*** Photomicrographs of pyramidal neurons in a living brain slice from a cFos-GFP rat sacrificed after the final EXT session (d28). Cells were visualized in the electrophysiology recording chamber by contrast (Dodt) optics, and over-layed with pseudocolor fluorescence to show a GFP-positive (GFP+; open arrowhead) and a GFP-negative (GFP ^―^; closed arrowhead) nucleus. Star indicates the recording electrode. All subjects in Figure 7 were sacrificed immediately following their final 90 min behavioral session. ***F,G*** Comparison of responses from GFP^+^ and GFP ^―^ cells from coca-SA (red) and saline-Y (blue) treated rats following EXT (F) or cue-reinstatement (G). Closed and open symbols denote responses in GFP+ and GFP ^―^ cells, respectively. Measurements included average firing (*inset*, I-O curves), individual firing (scatter plots of responses at 1 nA) and AHPs (scatter plots). Where possible, equal numbers of GFP^+^/GFP ^―^ cells in animals were sampled. Note the elevated firing and reduced AHPs only in a subset of GFP+ cells from coca-SA rats, but not from sal-Y rats. *Italicized* numbers in B indicate the number of animals, whereas those in C, F and G the number of cells/animal. Sidak multiple comparison post-test: ***p≤0.0001 comparing I/O (*insets*) after extinction or cue for cocaine-SA GFP ^―^ vs GFP^+^; ** p ≤ 0.01 comparing AHPs after extinction for GFP^+^ coca-SA vs GFP^+^ sal-Y; *p ≤0.05 comparing AHPs after cue for GFP^+^ coca-SA and GFP+ sal-Y.

### Pharmacological compounds

Cocaine hydrochloride was obtained through the Drug Supply Program of the National Institute on Drug Abuse. Dimethyl sulfoxide and D-glucose were purchased from Sigma (Sigma-Aldrich Corp., St. Louis, USA). All salts used in the preparation of physiological buffers were purchased from Fisher chemicals. Dopamine, XE-991, Kynurenic acid and MK-801 maleate (Dizocilpine) were purchased from Abcam. Retigabine was purchased from Axon Medchem.

### PFC Slice Preparation

Rats were killed by rapid decapitation, the brain was removed, and a tissue block containing the PL-PFC was placed in ice-cold and oxygenated artificial cerebrospinal fluid (aCSF) (in mM: 126 NaCl, 2.5 KCl, 1.2 MgCl_2_, 1.4 NaH_2_PO_4_, 25 NaHCO_3_, 11 D-glucose, 0.4 ascorbate) supplemented with MK-801 maleate (10 μM) and Kynurenic acid (2.6 mM) (Riegel and Williams, 2008; Williams et al., 2014). Thick sections (∼200 μM) containing the PL-PFC region (Figure 1*B*) were prepared using a vibrating microtome (Leica, Weltzar, Germany) and maintained in a vial containing MK-801 maleate (10 μM) dissolved in oxygenated aCSF in a 32°C water bath until recording. During recording, sections were perfused at a flow-rate of 2 ml/min with oxygenated aCSF at 33°C C. The prelimbic layer V/VI was identified initially using a 5X objective, and later magnified for patching with a 63X (0.9 N/A) water immersion lens fitted to an upright microscope (Olympus BX 51WI) equipped with gradient contrast infrared optics. GFP was visualized by fluorescence using a purpose-built LED system comprised of parts from Thorlabs.

### Electrophysiological procedures

Whole-cell recordings were performed using multiclamp 700B amplifiers (Molecular Devices). Neurons were voltage clamped at -70 mV using 1.5-2.5 MΩ glass microelectrodes filled with internal solution (in mM: 115 K-methylsulfate, 20 NaCl, 1.5 MgCl_2_, 2.5 HEPES, 2 ATP, 0.3 GTP, and 0.1 EGTA; pH 7.3; 265–270 mOsm). Series resistance (<10 mΩ) was compensated at 80%, and only cells with <15% series resistance change after drug application were included in the analysis. Recordings were collected online using AxoGraph X (AxoGraph Scientific) and digitized at 2–5 kHz. Layer V pyramidal cells were identified by location, morphology, and electrophysiological criteria including their characteristic hyperpolarized resting membrane potential and low input resistance (Gulledge and Jaffe, 1998; Ceci et al., 1999; Rosenkranz and Grace, 2002; Song et al., 2015). Passive membrane properties in these cells were consistent with published results (Gulledge and Jaffe, 1998; Ceci et al., 1999; Rosenkranz and Grace, 2002; Song et al., 2015) and did not vary between any of the treatment groups at any phase of behavioral training (average all groups across treatments: RMP: -67.2 ± 0.7 mV, *F*_(11, 610)_ = 1.084, p=0.3715; Rn: 58±2.0 MΩ, *F*_(11, 747)_ = 1.306, p=0.2160; Cm: 30.8 ± 1.0 pF, *F* _(11, 537)_ = 1.529, p=0.1172). Brain slices prepared at the culmination of the behavioral assay (Figure 1*B*) were used for current clamp (*I*-clamp) recordings in Experiment 3 (Figures 5 and 7). Experiments 1, 2 and 4 contain both *I*-clamp and voltage clamp (*V*-clamp) recordings (Figures 3, 4 and 6).

For *I*-clamp experiments the bridge was balanced routinely. Using published criteria, input-output (I-O) curves of action potential firing were generated by injecting a series of eight 0.8 s current steps, 1s interval (in pA: 100, 200, 300, 400, 500, 600, 800, 1000) and counting the numbers of action potentials (APs) evoked during the last 500 ms in each step epoch (Figure 1*B*) (Madison and Nicoll, 1984; Storm, 1990; Faber et al., 2001; Buchta et al., 2017). All firing traces shown represent the response to 1000pA current injections. Spike frequency adaptation (SFA) or accommodation is defined as the slowing of neuronal firing during a depolarizing current step (Madison and Nicoll, 1982; Lancaster and Nicoll, 1987; Aiken et al., 1995; Shah et al., 2006). Based on published studies, we identified SFA positive (SFA^+^) neurons as those showing 5 or fewer spikes during the final 500 ms of the 1 nA current injection (Madison and Nicoll, 1982; Lancaster and Nicoll, 1987; Aiken et al., 1995; Shah et al., 2006). AHPs were measured following calibration of a 60 ms current injection step to yield a burst of five APs from a holding potential of -70 mV (Figure 1*B*) (Coulter et al., 1989; Moyer et al., 2000; Gu et al., 2005). Because there are no clear criteria for separation of slow and medium AHP, we calculated total AHPs as the area below baseline following the 5th AP and return of the membrane potential to baseline values (Coulter et al., 1989; Moyer et al., 2000; Gu et al., 2005).

For voltage clamp experiments (Figure 1*C*), cells were first sorted as spike frequency adapting SFA(+) or not SFA(^―^) by a single 1000 pA / 800 ms current injection in *I*-clamp. After switching to *V*-clamp, 200nM tetrodotoxin (TTX) was continuously applied to the bath perfusate. After 5 min, outward currents were generated in response to a series of long (800 ms) depolarizing steps (-60 mV to -20 mV, 10 mV increment). Because Kv7 conductance is a component of this net outward current (and not the total current), the outward currents were measured again after perfusion of the Kv7 channel antagonist XE-991 (20 μM, 10 min) (Figure 1*C*). The long, slow *I*_Kv7_ activation kinetics and sensitivity to XE-991 have been characterized in previous reports (Delmas and Brown, 2005; Lawrence et al., 2006; Brown and Passmore, 2009). Consistent with published results, *I*_Kv7_ was voltage-dependent, outwardly rectifying, and active positive to -50 mV (Delmas and Brown, 2005; Lawrence et al., 2006; Brown and Passmore, 2009). The *I*_Kv7_ was measured by subtracting the outward currents generated in presence of XE-991 from the baseline (pre-drug) (Figure 1*C*) (Wang et al., 1998; Huang and Trussell, 2011). When the effects of dopamine or retigabine on Kv7 channel function was measured, cells were first sorted as SFA+/ ―, then voltage clamped and injected with a series of five long (1 s) depolarizing steps (-70 mV to -20 mV, 10 mV increment) in the presence of 200 nM TTX (Figure 1*C*). Outward current was measured 5 minutes after perfusion with 10 μM dopamine or retigabine 20 μM, and again 10 minutes after perfusion with dopamine + 20 μM XE-991 or retigabine + 20 μM XE-991 (Figure 1*C*). The amplitude of the Kv7 current before and after dopamine or retigabine treatment was computed by subtracting the final set of currents from the first and second sets of currents, respectively (Figure 1*C*). Recovery from dopamine or retigabine was not regularly examined, as full washout can exceed 25 min, a duration of time which negatively impacts the quality of patch clamp recordings in older animals that have undergone operant training (Yang and Seamans, 1996; Zheng et al., 1999; Henze et al., 2000; Gorelova et al., 2002). The 10 μM dopamine concentration for study was selected for its relevance to earlier studies using 10-30 μM (Malenka and Nicoll, 1986).

### Fos Immunohistochemistry

Immunohistochemical methods for verification of the c-fos-GFP rats were adapted from previous studies (Taniguchi et al., 2017). Fluorescence labeling was used to quantify the co-localization of the intrinsic GFP signal (c-fos) and the immediate early gene’s protein product (Fos) between different treatment groups of the self-administration paradigm in layer V of the PFC. Sections were washed 3x7 min in Phosphate-Buffered Saline (PBS), pH 7.4, and incubated for 1 hr in blocking solution (3% BSA, 0.3% Triton-X 100, 0.2% Tween, 3% normal donkey serum (NDS) in PBS). Sections were then incubated overnight at 4°;C with anti-Fos Goat primary antibody (sc-52-G, Lot B2513, Santa Cruz Biotechnology) diluted 1:1000 in blocking solution and anti-GFP Chicken (GFP-1020, Aves Labs Inc., Lot 0316FP11). Sections were washed in PBS 3x 7 min and incubated for 1.5 hr at room temperature with Donkey anti-chicken 488 (703-545-155, Jackson Immuno Research, Lot #: 119192) 1:200 and Donkey anti-goat Cy3 (705-165-147, Jackson Immuno Research, Lot # 132480) 1:200. Sections were washed, dehydrated, and mounted using ProGold Anti-Fade+ DAPI.

### c-Fos GFP and Fos Co-localization Quantification

Ninety minutes after the start of the final day of self-administration of cocaine or yoked-saline or the first day of extinction training, rats were deeply anesthetized with isofluorane as described elsewhere (Cifani et al., 2012), and transcardially perfused (150 mL of saline and 150 mL of 4% PFA in PBS). Whole brain specimens were postfixed in 4% PFA for 24 hrs, and transferred into PBS for 24 hrs until immunohistochemistry procedures were conducted to confirm the c-Fos GFP transgene as described elsewhere (Cifani et al., 2012). The total c-Fos GFP/anti-Fos cell counts, the Pearson’s Correlation Coefficients, and the Mander’s Overlap Coefficients describing the co-expression of antibody staining for Fos and c-Fos GFP were obtained to verify the efficacy of the c-Fos GFP transgene (Figure 7). A confocal microscope (Zeiss LSM 510) was used to image paraformaldehyde fixed brain sections. Fluorescent images of layer V PFC were taken using both Leica and NIS Elements Software. Quantification of co-localization of the GFP expression (indicative of c-fos) and the fos immunofluorescence was conducted using the Nikon NIS Elements Software on 10x bilateral images of layer V PFC.

### Western blotting

Samples of the PL-PFC were lysed by sonication in RIPA buffer (50mM Tris-HCl, pH 7.4, 50mM NaCl, 2mM MgCl_2_, 1% [vol/vol] NP40, 0.5% [mass/vol] sodium deoxycholate, and 0.1% [mass/vol] sodium dodecyl sulfate), protease (Cat# B14002; Biotool) and phosphatase inhibitors (Cat# B15002, Biotool), and BitNuclease (Cat# B16002, Biotool). After sonication, lysates were clarified by centrifugation (15000g, 10 min, 4°;C), then protein concentrations were determined using the BCA protein assay (Cat# PI23225, Thermo Fisher Scientific). Approximately 15 μg of total proteins were loaded on an SDS-PAGE and then transferred to polyvinylidene difluoride membranes (0.45 μm) and blocked at room temperature for 1 hr in TBST (50 mM Tris-HCl, pH 7.4, 150 mM NaCl, 0.1 % Tween 20), 5% non-fat dry milk. Primary antibodies used for probing KCNQ2 (Cat# APC-050, Alomone, Jerusalem, Israel), KCNQ3 (Cat# APC-051, Alomone), KCNQ5 (Cat# APC-155, Alomone) and Actin (Cat# A2066, Sigma) were diluted in TBST with 5% bovine serum albumin. Immunoblots were revealed by enhanced luminescence (WBKLS0500, Millipore) before exposure to a photographic film. Films were scanned, digitized, and quantified using Un-Scan-It gel version 6.1 scanning software (Silk Scientific Inc).

Experimental Design and Statistical Analysis. All data are reported as mean ± SEM, and statistical analyses were performed utilizing GraphPad Prism. Electrophysiological data were analyzed using Axograph X. The effects of bath applied dopamine were evaluated using paired-*t* tests. AHP differences expressed as area under the curve were evaluated using one-way measures analyses of variance (ANOVAs) or unpaired *t*-tests. Two-way mixed repeated measures analyses of variance (ANOVAs) were used to evaluate the effects of treatment (cocaine, saline, sucrose), excitability (firing or current amplitude) and time (repeated steps of current or membrane potential). All statistical evaluations of Kv7 channel I-V curves were conducted at data points positive to –50 mV, when channels are open and not inactivated (Lawrence et al., 2006; Huang and Trussell, 2011; Battefeld et al., 2014). The between-subject variable was treatment (cocaine-SA, coca-Y, sal-Y, sucrose-SA) and the repeated measure was current (for action potential firing) or membrane potential (for current amplitude). Significant interactions (treatment × current or treatment × membrane potential) prompted separate two-way repeated measures ANOVA for each treatment. Current and action potential responses of control and treated rats were typically compared at the maximal difference (1 nA for action potential firing or - 20 mV for membrane currents). The effects of bath applied drug (retigabine, XE-991, or dopamine) were evaluated using two-way repeated measures ANOVAs, with the factors of current (or membrane potential), drug (retigabine, XE-991, or dopamine) and treatment (or SFA). All pairwise comparisons were conducted using Sidak multiple comparisons test. For graphical purposes and clarity, data are often presented separately for cocaine and saline treatment groups. Data represent the mean ± SEM, with a α set at *P* < .05.

## Results

This study addressed three interrelated questions: 1) What are the long-lasting alterations in PL-PFC excitability that occur following cocaine self-administration (withdrawal and extinction)? 2) What are the acute alterations in excitability that occur immediately following cue-induced reinstatement of cocaine seeking? 3) Do the observed alterations contribute to the cue-induced reinstatement of cocaine seeking?

To address these questions, we employed the self-administration model of relapse to drug-seeking (Shaham et al., 2003). Rats were trained to self-administer cocaine (0.20 mg/kg/infusion) for 14 days in operant chambers where drug infusions were paired with light/tone cues (Figure 1*A*). All rats showed stable cocaine self-administration (coca-SA) responding (Figure 2*A*). Parallel groups of rats received yoked infusions of saline or yoked infusions of cocaine (Figure 2*A*). No differences in the total number of drug or saline infusions were found between the three groups (one-way ANOVA, *F*_(2, 70)_ = 1.819, p = 0.1697). As expected, lever pressing in the three treatment groups differed (Figure 2*A*; two-way ANOVA with repeated measures over time, treatment *F*_(2,609)_ = 91.33, p < 0.0001; time *F*_(14,609)_ = 0.3788, p = 0.9806; interaction *F*_(28,609)_ = 0.8388, p = 0.7060). Coca-SA rats pressed the active lever significantly more than sal-Y or cocaine-Y controls (Figure 2*A*; coca-SA vs sal-Y *t*_(609)_ = 12.17 p < 0.0001; coca-SA vs coca-Y, *t*_(609)_ = 9.205 p < 0.0001; sal-Y vs coca-Y *t*_(609)_ = 0.4642 p = 0.9544). Self-administration training was followed by 14 to 19 days of extinction (Ext) where neither cues nor drug was available. As previously reported in coca-SA treated rats, the first day of extinction (Figure *2A*; vertical dashed line) was associated with a burst in lever pressing (Lerman and Iwata, 1995). As expected, rats trained for coca-SA showed stable context extinction of responding on the active levers (Figure 2*A*).

Rats from the coca-SA and sal-Y groups also underwent testing for cued-reinstatement (Figure 2*B)*. Relapse to cocaine seeking was modeled in a session where cocaine-associated cues were presented, but lever pressing did not result in drug delivery (Kalivas and McFarland, 2003; McLaughlin and See, 2003). Responses to reinstatement testing differed between the sal-Y and cocaine-SA groups (two-way ANOVA, treatment *F*_(1,33)_ = 95.12, p < 0.0001; time *F*_(1,33)_ = 105.6, p < 0.0001; interaction *F*_(1,33)_ = 61.57, p < 0.0001). Lever pressing in the coca-SA group increased above that observed during the last day of extinction training (Figure 2*B*; *t*_(33)_ = 13.8, p < 0.0001). As predicted, rats in the sal-Y group did not show increased lever pressing in response to light and tone cues (Figure 2*B*; *t*_(33)_ = 1.612, p = 0.2194).

### Active (but not passive) exposure to cocaine suppresses dopamine-sensitive inhibition

Experiment 1: To determine if cocaine self-administration causes acute changes in PL-PFC excitability, we sacrificed animals 24 hours after their last self-administration training session. In acute brain slices, we recorded Layer V pyramidal neurons in the PFC in *I*-clamp to measure action potential firing to depolarizing current injections steps (0.8 s, 0.1-1 nA). We observed obvious differences in the firing responses between coca-SA, coca-Y, and sal-Y treatment groups during the depolarizing current steps (Figure 3*B1*, *inset*; two-way ANOVA with repeated measures over current, treatment *F*_(2,76)_ = 34.89, p<0.0001, current *F*_(7,532)_ = 62.94, p<0.0001, interaction *F*_(14,532)_ = 19.90, p<0.0001). A post hoc analysis showed no significant firing differences between the two yoked-treatment groups (p = 0.6400), but a significant increase in coca-SA responses relative to the coca-Y group (p < 0.0001), which was surprising given that the two groups received similar amounts of cocaine (Figure 2*A*). As prior reports show heterogeneous subpopulations of cortical neurons (Cruz et al., 2013b; 2015), we used scatter plots to compare the relative distribution of individual firing responses at the 1 nA current step. We categorized cell activity according to the degree of spike-frequency adaptation (SFA), whereby SFA positive (SFA^+^) neurons were classified as those showing 5 or fewer spikes during the final 500 ms of the 1 nA current injection (Figure 3*B1*). In most coca-SA neurons (81%), training experience elevated firing, i.e. above the SFA threshold of 5 spikes (Figure 3*B1*). By contrast, most firing responses in sal-Y (9%) or non-contingent coca-Y (21%) groups displayed robust SFA (i.e. SFA^+^) (Figure 3*B1*). The SFA is modulated by the after-hyperpolarization (AHP) that follows a depolarization step (Madison and Nicoll,1986). AHPs were measured following calibration of a 60 ms current injection step to yield a burst of five APs from a holding potential of -70 mV (Figure 1*B*) and measured the total AHP area (Coulter et al., 1989; Moyer et al., 2000; Gu et al., 2005). As expected, after training, the AHP size decreased in the coca-SA group, but not the yoked groups (Figure 3*B2*; one-way ANOVA: treatment *F*_(2,41)_ = 13.92, p<0.0001; post hoc test, coca-SA vs: sal-Y p < 0.0001; coca-Y p = 0.0044). These data suggest learning to lever press for cocaine and not the mere exposure to cocaine itself increases neuronal excitability.

In other cortical regions, activation of Kv7 voltage-gated K^+^ channels localized to the axon hillock suppresses neuronal output by enhancing SFA (Delmas and Brown, 2005; Lawrence et al., 2006; Brown and Passmore, 2009). To determine if coca-SA experience reduced this form of inhibition in PL-PFC cells, we sampled a range of membrane potentials in *V-*clamp (-60 to -20 mV) to measure Kv7 channel currents (*I*_Kv7_; *inset* Figure 3*C1*). We observed voltage-dependent *I*_Kv7_ activated above -50 mV (Figure 3*C1*). As with the previous firing measurements, the amplitude of *I*_Kv7_ also differed significantly between the coca-SA and sal-Y treatments (Figure 3*C1*; two-way ANOVA with repeated measures of potential, treatment *F*_(1,8)_ = 5.973, p<0.0403, potential *F*_(2,16)_ =26.67, p<0.0001, interaction *F*_(4,32)_ = 5.41, p<0.0019). When considering all cells sampled from the coca-SA and Y-sal groups, the elevated basal level of firing at 1 nA correlated with the reduction in *I*_Kv7_ amplitudes (Figure 3*C2*; R^2^ = 0.479). Thus, at 24 hrs following the 2 weeks of cocaine self-administration, the loss of SFA may be due to functional suppression in Kv7 channel mediated inhibition.

Our previous published work showed that bath application of dopamine or chemo/optogenetic activation of VTA dopamine terminals suppressed SFA and AHPs in PL-PFC pyramidal cells (Buchta et al., 2017). Since repeated activation of cAMP-coupled GPCRs can sensitize adenylate cyclase to subsequently activate downstream signaling cascades (Johnston and Watts, 2003), we wondered if dopamine suppresses SFA by closure of Kv7 channels and whether the repeated coca-SA impacted this function. To evaluate this, we sampled SFA, AHP and *I*_Kv7_ before and during application of 10 μM dopamine in the sal-Y, coca-Y and coca-SA group (Figure 3*D* and *E*).

At the 1 nA current injection step, all sampled PL-PFC cells in the coca-SA group lacked SFA, whereas all cells in the sal-Y and coca-Y groups displayed SFA (Figure 3*D1*). In SFA^+^ cells from both the sal-Y and coca-Y groups, dopamine increased firing responses and decreased AHPs relative to the pre-dopamine levels (Figure 3*D1*; paired *t*-test firing: sal-Y/SFA^+^ *t*_(8)_ = 9.06, p<0.0001; coca-Y/SFA^+^ *t*_(8)_ = 14.06, p<0.0001; Figure 3*D2*, paired *t*-test AHPs: sal-Y *t*_(6)_ = 5.581, p<0.0014; coca-Y *t*_(9)_ = 8.199, p<0.0001). By contrast, firing and AHP responses in the SFA ^―^ cells of the coca-SA group were not measurably changed by dopamine figure 3*D1*; paired *t*-test: firing, *t*_(9)_ = 1.262, p=0.2385; Figure 3*D2*: AHP: *t*_(9)_ = 1.333, p=0.2153). Given the diminished sensitivity to dopamine in the firing and AHP experiments in the coca-SA group, we examined the dopamine action on evoked *I*_Kv7_ currents in cells of the cocaine-SA group, all of which were SFA ^―^ (Figure 3*E*). *I*_Kv7_ responses to dopamine in coca-SA and sal-Y treatments differed significantly (two-way ANOVA with repeated measures of potential, treatment *F*_(3,38)_ = 5.902, p=0.0021, potential *F*_(2,76)_ =65.03, p<0.0001, interaction *F*_(6,76)_= 8.949, p<0.0001). To understand the nature of the interaction, we deconstructed individual treatment responses at -20 mV, a potential where *I*_Kv7_ was maximal. In support of our results above in Figure 3*C1*, follow up comparisons between treatments showed significantly reduced *I*_Kv7_ amplitudes under baseline (pre-dopamine) conditions in the cocaine-SA treatment group relative to yoked-saline controls (p < 0.0001). Furthermore, dopamine reduced the *I*_Kv7_ in the yoked-saline group (p < 0.0001), but not in the cocaine-SA group (p = 0.6544) (Figure 3*E)*. In summary, the active self-administration (but not passive infusion) of cocaine was associated with a reduction in intrinsic inhibition, reduced sensitivity to dopamine, and suppression of inhibitory Kv7 ion channel currents.

### Dopamine-sensitive inhibition is suppressed after 14 days extinction training

Experiment 2: To determine if the changes in PL-PFC excitability persisted after 14d of extinction conditioning, rats were sacrificed 15 min following the final extinction session (Figure 4*A*). We excluded the coca-Y group from further analysis since the previous experiment already showed that intrinsic inhibition was not altered by passive administration of cocaine, only by active self-administration of cocaine. We found that after extinction training, average firing responses were different between the coca-SA and sal-Y control treatments (Figure 4*B1*; two-way ANOVA with repeated measures of current, treatment *F*_(1,66)_ = 5.309, p = 0.0244; current *F*_(7,462)_ = 13.73, p<0.0001; interaction *F*_(7,462)_ = 4.694, p < 0.0001). Firing responses at 1 nA current step were elevated in the coca-SA group relative to sal-Y controls (*t*_(528)_ = 4.372, p = 0.0002). Roughly half (57%) of coca-SA cells showed high firing (SFA ^―^ responses), whereas only a minority (11%) of sal-Y control cells were SFA^―^Compared to training alone (without extinction; Figure 3), there was a slight reduction in SFA ^―^ cells (from 81% to 57%) in the coca-SA group. Examination of individual cells showed evenly distributed responses in sal-Y treatments, but heterogeneous responses in the coca-SA treatments (Figure 4*B2*), suggesting that only a subset of cells are persistently hyperactive after extinction training. AHP measurements following extinction showed similar changes; in the coca-SA treatment group the mean value declined, but individual cells tended to cluster at higher or lower responses (Figure 4*B3*; unpaired t test; *t*_(80)_ = 3.898, p = 0.0002). Together, these data suggest that following extinction, a sub-population of SFA ^―^ coca-SA cells remain hypersensitive to depolarizing inputs.

To examine the involvement of Kv7 channels, first we measured the effects of the channel antagonist XE-991 (20 μM, 10 min) on firing frequency and second measured the *I*_Kv7_ in voltage-clamp in cells from the sal-Y group and the population of SFA ^―^ cells from coca-SA rats immediately after the final extinction session. First, responses to XE-991 differed in sal-Y SFA^+^ cells and the subpopulation of SFA ^―^ cells from the coca-SA rats (two-way ANOVA with repeated measures of current, treatment *F*_(3,30)_ = 17.89, p < 0.0001; current *F*_(7,210)_ = 94.48, p<0.0001; interaction *F*_(21,210)_ = 10.23, p < 0.0001; data not shown). A subsequent analysis within the sal-Y group showed XE-991 increased firing across the full range of current steps, culminating in a 10-fold potentiation at the 1 nA current step (two-way ANOVA with repeated measures of current, XE991 *F*_(1,112)_ = 42.42, p=0.0001; current *F*_(7,112)_ = 10.06, p<0.0001, interaction *F*_(7,112)_ = 6.971, p<0.0001). By contrast, application of XE-991 did not alter the input-output curve for SFA ^―^ cells within the coca-SA group (two-way ANOVA with repeated measures of current, XE-991 *F*_(1,16)_ = 0.5168, p=0.4826; current *F*_(7,112)_ = 95.73, p<0.0001; interaction *F*_(7,112)_ = 0.8562, p = 0.5435). A follow up analysis of baseline (pre-XE-991) conditions comparing sal-Y and coca-SA treatments at different current steps showed elevated action potential firing in the coca-SA (SFA ^―^ cells) group, indicating firing may already be maximal and therefore less prone to further potentiation by blockade of Kv7 channels (two-way ANOVA with repeated measures of current, treatment *F*_(1,15)_= 42.82, p=0.0001; current *F*_(7,105)_ = 30.83, p<0.0001; interaction *F*_(7,105)_ = 23.46, p<0.0001). Taken together, these data indicate that after extinction from cocaine self-administration, a population of PL-PFC cells show enhanced firing responses under baseline conditions and a related diminished sensitivity to Kv7 channel block.

Second, to corroborate these results, we measured *I*_Kv7_ responses (Figure 4*C1*, *inset*) in all cells across membrane potentials. We found significant differences between the 3 groups (sal-Y/SFA^+^, coca-SA/SFA^+^, coca-SA/SFA ^―^) (Figure 4*C1*; two-way ANOVA with repeated measures of potential, group/SFA *F*_(2,29)_ =8.669, p = 0.0015; potential *F*_(2,48)_ = 58.94, p<0.0001; interaction *F*_(4,48)_ = 4.344, p = 0.0044). At -20 mV, *I*_Kv7_ responses in SFA ^―^ cells in the coca-SA group were reduced relative to the SFA^+^ cells in the sal-Y group (Figure 4*C1*; coca-SA/SFA^+^ vs coca-SA/SFA ^―^: *t*_(72)_ = 3.817, p = 0.0008; sal-Y/SFA^+^ vs coca-SA/SFA ^―^: *t*_(72)_ = 5.634 p <0.0001; sal-Y vs coca-SA/SFA^+^: *t*_(72)_ = 2.211, p = 0.0878). Similar to findings at 24hr after cocaine SA, the basal level of firing at 1 nA correlated with *I*_Kv7_ amplitudes (Figure 4*C2*; ext R^2^ = 0.370). These results highlight an enduring functional shift in inhibition in a population of SFA ^―^ cells in the coca-SA group.

Given that dopamine regulates SFA in the PFC, we examined whether this regulation is altered following extinction (Figure 4*D)*. In SFA^+^ cells of either sal-Y or coca-SA rats, bath application of dopamine (10 μM, 10 min) produced robust changes in SFA and AHPs (paired *t*-test; Figure 4*D1*, sal-Y/SFA^+^: *t*_(8)_ = 4.664, p < 0.0001; coca-SA/SFA+ *t*_(5)_ = 6.677, p = 0.0011; Figure 4*D2*, sal-Y/SFA^+^ *t*_(9)_ = 6.335, p < 0.0001; coca-SA/SFA+ *t*_(3)_ = 6.976, p = 0.0060). In contrast, in SFA ^―^ cells (only observed in the coca-SA group), there was no measurable dopamine-induced change in firing or AHPs (Figure 4*D1*; paired *t*-test firing: t_(7)_: = 0.1573, p = 0.8795; Figure 4*D2*, paired *t*-test AHP: t_(6)_ = 0.9185, p = 0.4005). Correspondingly, responses of inhibitory Kv7 currents to dopamine also differed between treatment groups (Figure 4*E*; two-way ANOVA with repeated measures of potential, treatment *F*_(5,42)_ = 5.025, p = 0.0011; potential *F*_(2,84)_ = 75.74, p = 0.0001; interaction: *F*_(10,84)_ = 7.968, p < 0.0001). This difference could be explained, in a subsequent analysis, by the finding that most cells in the sal-Y group were SFA^+^. However, in both the sal-Y/SFA^+^ and coca-SA/SFA^+^ groups, dopamine reduced *I*_Kv7_ amplitudes (two-way ANOVA with repeated measures of potential: sal-Y/SFA^+^, dopamine *F*_(1,16)_ = 6.534, p = 0.0201; potential *F*_(2,32)_ = 28.37, p = 0.0001; interaction: *F*_(2,32)_ = 6.185, p < 0.0054; coca-SA/SFA^+^, dopamine *F*_(1,12)_ = 5.646, p = 0.0350; potential *F*_(2,24)_ = 31.5, p = 0.0001; interaction: *F*_(2,24)_ = 7.927, p = 0.0023). In contrast, dopamine did not alter *I*_Kv7_ amplitudes in coca-SA/SFA ^―^ cells (Figure 4*E*; two way ANOVA with repeated measures of potential, dopamine *F*_(1,14)_ = 3.695, p = 0.0752; potential *F*_(2,28)_ = 19.44, p < 0.0001; interaction: *F*_(2,28)_ = 9.973, p < 0.0005). A final comparison of the baseline (pre-dopamine, -20 mV) conditions between the three groups showed significantly reduced *I*_Kv7_ amplitudes in the coca-SA/SFA ^―^ group relative to coca-SA/SFA^+^ cells (Figure 4*E*; one-way ANOVA *F*_(2,21)_ = 3.653, p = 0.0435; post hoc sal-Y/SFA^+^ versus: coca-SA/SFA^+^, *t*_(21)_=1.578, p =0.3404, coca-SA/SFA ^―^, *t*_(21)_= 1.936, p =0.1865; coca-SA/SFA^+^ versus coca-SA/SFA ^―^: *t*_(21)_=3.354, p = 0.0090). This suggests the baseline amplitude of *I*_Kv7_ in coca-SA/SFA ^―^ cells was already reduced prior to bath application of dopamine. Together, these data indicate after 2 weeks of cocaine self-administration, a population of SFA ^―^ cells experience a reduction in intrinsic inhibition and therefore reduced sensitivity to dopamine regulation, and this relationship persists even after the behavioral extinction of drug-paired associations.

### Loss of inhibition is independent of extinction training and does not persist after extinction from sucrose-SA

Experiment 3: Typically, reductions in intrinsic inhibition are transient, learning-specific, and behaviorally appropriate (Oh and Disterhoft, 2015). To determine if extinction learning is required for the enduring reduction in intrinsic inhibition in the coca-SA/SFA ^―^ population, we sampled cells from coca-SA animals that did not undergo extinction training, but instead underwent 14d of homecage withdrawal (Figure 5*A*). Like the firing responses in extinguished animals (Figure 4*B*), average firing responses in about half of cells in the withdrawal group exceeded the maximal limit for SFA, and the individual responses for firing and AHPs clustered at high and low values (Figure 5*A*). This indicates that intrinsic inhibition is also modified in cocaine-SA animals that did not experience extinction training. In addition, we sacrificed a group of naÏve animals without any prior experience of handling and surgery. Cells from naÏve rats displayed firing and AHP responses (Figure 5*B*) qualitatively identical to those observed after sal-Y treatment (Figure 4*B*). This indicates that the cocaine-SA induced adaption in intrinsic inhibition in the PFC did not require extinction training or the handling associated with the surgical procedures.

Self-administration of sucrose can modify some of the same brain reward circuitry stimulated by drugs of abuse (McGlinchey et al., 2016). We next determined if sucrose, a nondrug reinforcer, produced adaptations in intrinsic inhibition in the PFC similar to that observed after cocaine self-administration. Rats trained to self-administer sucrose pellets paired with light/tone cues (Figure 5*C1*) for 10-14 days showed stable lever-pressing (Figure 5*C2*). During extinction conditions (Ext, 14 to 19 days) when neither cues nor sucrose was available, lever-press responding diminished as expected (Figure 5*C2*). To evaluate firing and AHP responses, rats were sacrificed either at 24 hr after the final self-administration training session (Training; d15) or immediately after the final extinction (Ext; d29). Firing responses in the sucrose-SA rats after training and extinction differed (Figure 5*C3*; two-way ANOVA with repeated measures of current, treatment *F*_(1,45)_ = 5.525, p = 0.0232; current *F*_(7,315)_ = 20.97, p < 0.0001; interaction *F*_(7,315)_ = 4.921, p < 0.0001). After training, roughly half of the sampled sucrose-SA cells showed elevated firing (SFA ^―^, 56%) (Figure 5*C4*), a much lower proportion than we observed after cocaine-SA training (Figure *3B1*, 81%). In contrast, following extinction, few cells in the sucrose-SA group showed elevated firing (Figure 5*C4*, SFA ^―^, 14%). In fact, sucrose-SA responses after extinction more resembled those described previously in the sal-Y with extinction (Figure 4*B*, 11%) or in naÏve rats (Figure 5*B*, 5%), and did not resemble responses in the cocaine-SA extinction group (Figure 4*B2*, 57%). AHPs in the sucrose-SA rats after training and extinction also differed (Figure 5*C5*; unpaired t test; *t* _(41)_ = 6.876, p < 0.0001). AHP responses in the sucrose-SA with extinction group resembled responses in the sal-Y with extinction (Figure 4*B*) or in naÏve rats (Figure 5*B*, 5%), and diverged from AHP responses in the cocaine-SA extinction group (Figure 4*B3*). Taken together, this indicates that training to self-administer drug (cocaine) and nondrug (sucrose) reinforcers produce qualitatively similar reductions in intrinsic inhibition early on, but for drug reinforcers these modifications are persistent even after withdrawal or extinction. These data are consistent with previous studies reporting that drug and nondrug reinforcers produce different cellular adaptations in PL-PFC neurons (Whitaker et al., 2017).

### Stabilization of Kv7 mediated inhibition suppresses cued-reinstatement

Experiment 4: To determine if loss of intrinsic inhibition contributes to relapse and to what extent stabilization of inhibitory Kv7 channels would reduce relapse, we measured responses in rats 15 min after a 2 hr cue-reinstatement session (Figure 6*A*). Following cued-reinstatement testing (Figure 2*B*), the coca-SA group showed robust differences in firing from sal-Y controls (Figure 6*B1*; two-way ANOVA: treatment *F*_(1,43)_ = 34.22, p<0.0001, current *F*_(7,301)_ = 36.09, p<0.0001, interaction *F*_(7,301)_ = 23.16, p<0.0001). A post hoc analysis showed elevated firing (SFA ^―^ responses) in the coca-SA group (1 nA current step, *t*_(344)_ = 10.68, p < 0.0001). Whereas after cue testing most coca-SA cells (72%) showed this firing response, only a minority (11%) of sal-Y controls were SFA ^―^ (Figure 6*B2*). Although visual inspection of dot plots depicting individual cell AHP responses in coca-SA and sal-Y groups showed an apparent divergent distribution of responses in the coca-SA group (Figure 6*B3*), the difference in the average AHP between groups did not reach statistical significance (Figure 6*B3*; *t*-test: treatment *t*_(41)_ = 1.847, p = 0.0720). Thus, cued-reinstatement is associated with potentiated firing responses in SFA ^―^ cells from rats with extinguished behavioral responding for cocaine self-administration.

Because changes in SFA responses following reinstatement testing exceeded those following extinction, we determined if cued-reinstatement exposure also suppressed the function of Kv7 channels (Figure 6*C*). A comparison of *I*_Kv7_ function after cued-reinstatement testing showed obvious differences between the SFA^+^ cells (from both the sal-Y and coca-SA groups) and SFA ^―^ cells in the coca-SA group (Figure 6*C*; two-way ANOVA with repeated measures of potential, treatment *F*_(2,29)_ = 9.064, p=0.0006; potential *F*_(2,58)_ = 80.98, p<0.0001; and interaction: *F*_(4,58)_ = 7.272, p<0.0001). A post hoc analysis of responses at -20 mV indicated no difference in *I*_Kv7_ amplitude between SFA^+^ cells from sal-Y and coca-SA rats (Figure 6*C*; *t*_(87)_ = 0.8783, p=0.7673), but a significant reduction of *I*_Kv7_ amplitudes in the SFA ^―^ cell population (Figure 6*C*;coca-SA/SFA ^―^ vs: coca-SA/SFA^+^ *t*_(87)_ = 4.718, p<0.0001; sal-Y/SFA^+^ *t*_(87)_ = 6.166, p<0.0001). Thus, the function of Kv7 channels is suppressed in SFA ^―^ cells following reexposure to drug paired cues that reinstate drug seeking

To determine if stabilization of Kv7 channels with retigabine could restore SFA and *I*_Kv7_ in neurons from coca-SA experienced animals, two types of experiments were performed. In the first experiment, current clamp recordings were conducted in slices prepared at 15 min after a 2hr cue-test, in order to sample firing both before and after bath application of retigabine (20 μM, 10 min). Firing responses in the sal-Y and cocaine-SA treatments differed considerably at this time point (Figure 6*D*; two-way ANOVA: treatment *F*_(3,34)_ = 22.44, p=0.0001; current *F*_(7, 238)_ = 9.626, p=0.0001; interaction *F*_(21, 238)_ = 14.49, p < 0.0001). A post hoc analysis of the retigabine action measured at the 1 nA current step indicated no measurable retigabine effect on firing in SFA^+^ cells from sal-Y rats (*t*_(272)_ = 0.1590, p > 0.9999), but a robust retigabine mediated reduction of firing in SFA ^―^ cells from coca-SA rats (*t*_(272)_ = 14.2, p<0.0001). Presumably, the nominal response of controls to retigabine reflected strong preexisting intrinsic inhibition, whereas the pronounced retigabine action across the full range of current injections in SFA ^―^ cells in coca-SA rats reflected a preexisting state of weakened Kv7 channel mediated inhibition.

To test this notion, a second group of experiments determined if exposure to drug-paired cues reduced Kv7 channel inhibition and to what extent the reduction was rescued by retigabine. Following the cued-reinstatement test, *I*_Kv7_ currents were measured before and after bath application of retigabine (20 μM, 10 min). Retigabine responses in SFA^+^ cells from sal-Y controls and SFA ^―^ cells from coca-SA rats differed (Figure 6*E*;two-way ANOVA; treatment *F*_(3,32)_ = 5.503, p = 0.0037, potential *F*_(2,64)_ = 81.19; p < 0.0001, interaction *F*_(6, 64)_ = 3.559, p = 0.0042). Subsequent analysis showed retigabine increased the amplitude of *I*_Kv7_ in SFA ^―^ cells from coca-SA rats (*t*_(96)_ = 1.672, p = 0.0138), without significantly altering the *I*_Kv7_ responses in SFA^+^ cells from sal-Y rats (*t*_(96)_ = 1.672, p = 0.4609). This indicates that retigabine partially restored Kv7 channel inhibition to SFA ^―^ cells from coca-SA rats, but in contrast produced little measurable change in neurons with normal preexisting levels of intrinsic inhibition.

Cue-induced reinstatement of cocaine seeking is blocked by reversible pharmacological or chemo/optogenetic inactivation of cells in the PL-PFC or their terminals in the nucleus accumbens (Kalivas and McFarland, 2003; Gipson et al., 2013; 2014; Stefanik et al., 2016). It is possible that the drug-paired cues activated PL-PFC cells with reduced intrinsic inhibition, thus causing enhanced firing that promotes cocaine-seeking. To determine if stabilization of inhibitory PL-PFC Kv7 channels would reduce cued-reinstatement of cocaine seeking behavior, we used a counter balanced design using coca-SA rats that were extinguished from cocaine-seeking and microinjected retigabine or vehicle directly into PL-PFC region 10 minutes prior to reinstatement testing. Infusions altered the average lever-pressing behavior for drug associated cues (Figure 6*F*; two-way ANOVA, concentration *F*_(4,110)_ = 5.561; p = 0.0004, lever *F*_(1,110)_ = 56.64, p < 0.0001, interaction *F*_(4,_110) = 5.504, p = 0.0004). A post hoc comparison of pressing between the last extinction session and the cue session showed potentiated active-lever pressing in vehicle-injected animals (Figure 6*F*; *t*_(110)_ = 6.233, p<0.0001). The magnitude of the reinstatement was comparable to what we observed in our earlier experiments (Figure 2*B*) and consistent with previous reports (Venniro et al., 2016). Relative to vehicle infusions, retigabine dose-dependently reduced the reinstatement of active-lever pressing significantly at 300μM (Figure 6*G*; 3 μM *t*_(110)_ = 0.8641, p=0.9928; 30 μM *t*_(110)_ = 2.073, p=0.3387; 300 μM *t*_(110)_ = 3.036, p = 0.0295). A comparison of the total distance traveled by the same rats in a novel chamber locomotion test did not show significant differences caused by retigabine in any of the treatment groups (one-way ANOVA comparing vehicle, retigabine (30 μM), retigabine (300 μM): *F*_(2,33)_ = 0.5745, p=0.5685), suggesting the retigabine action on reinstatement was not due to a nonspecific reduction in motor behavior. The cannula placements for PL-PFC microinjections were verified histologically in all animals included in the analysis.

To determine if these cocaine experiences altered protein expression of Kv7 channel subunits, separate groups of sal-Y and coca-SA rats were sacrificed 15 min after the 2 hr reinstatement test for a Western blot analysis (Figure 6*G*). A comparison of the PL-PFC tissue-homogenates showed no differences in expression levels of the 3 principle subunits of the neuronal Kv7 family (Kv7.2, Kv7.3, and Kv7.5) (Figure 5*G*; two-way ANOVA, treatment *F*_(3,84)_ = 0.6641, p=0.5764; subunits: *F*_(2,84)_ = 0.3579, p=0.7002; interaction *F*_(6,84)_ = 7699, p = 0.5957). Taken together, these results suggest that rather than reducing Kv7 channel expression, the operant training for cocaine desensitizes the function of inhibitory Kv7 channels. Thus, by stabilizing the open state of Kv7 channels, retigabine reduces cue-induced reinstatement of cocaine seeking when injected into the PL-PFC.

### Reduced inhibition in cFos-GFP^+^, but not cFos-GFP^―^cells

Experiment 5: The experiments above used a generalized selection criterion of PL-PFC neurons based on location, morphology, and electrophysiological parameters to show that re-exposure to drug conditioned cues to reinstate drug seeking also rekindled the reduction in intrinsic inhibition that emerges after cocaine self-administration. This occurred in the majority of SFA ^―^ cells, despite the extinction of behavioral responding for cocaine. Because published studies have shown a causal link between the persistent expression of cocaine-related behaviors and activation of select ensembles of neurons following re-exposure to drug-associated cues and context (Koya et al., 2009; Bossert et al., 2011; Fanous et al., 2012; Bobadilla et al., 2017), we evaluated if the PL-PFC neurons with reduced inhibition represented a similar engram. Behaviorally relevant ensembles of cells have been identified in behaviorally activated cFos-GFP transgenic rats, where activation of the *c-fos* promoter drives expression of green fluorescent protein (*cFos*-GFP, Figure 7*A*) (Koya et al., 2012). Using the same line of transgenic animals, we determined if the expression of Fos paralleled the coca-SA modification in intrinsic inhibition.

Transgenic rats underwent self-administration training for cocaine or yoked infusion of saline followed by extinction (Figure 7*B*) and cued-reinstatement (Figure 7*C)*. No differences in the total number of drug or saline infusions were found between the group of transgenic rats and the wild-type rats examined above in Figure 2*A* (one-way ANOVA, *F*_(3, 90)_ = 0.3502, p = 0.7891). Similar to the above responses in wild-type rats (Figure 2*A*), transgenic rats trained to self-administer cocaine pressed the active lever significantly more than sal-Y controls (Figure 7*B*; two-way ANOVA with repeated measures over time, treatment *F*_(1,16)_ = 21.89, p=0.0003; time *F*_(13,208)_ = 1.221, p = 0.2661; interaction *F*_(13,208)_ = 1.963, p = 0.0253). Active-lever pressing for drug was extinguished over two weeks of extinction training (Figure 7*B*). The responses to reinstatement testing differed between the sal-Y and cocaine-SA groups (two-way ANOVA, treatment *F*_(1,25)_ = 34.56, p < 0.0001; time *F*_(1,25)_ = 35.38, p < 0.0001; interaction *F*_(1,25)_ = 38.10, p < 0.0001). Lever pressing in the transgenic coca-SA group increased above that observed during the last day of extinction training (Figure 7*C*; *t*_(25)_ = 8.649, p<0.0001). As predicted, rats in the transgenic sal-Y group did not show increased lever pressing in response to light and tone cues (Figure 7*C*; *t*_(25)_ = 0.1576, p = 0.9846).

To corroborate previous reports validating the transgene, one rat from each of the sal-Y and the coca-SA training groups received cardiac perfusion 15 min following the first extinction session (d15) for subsequent immunofluorescent evaluation. This time point is associated with a burst of lever pressing activity in coca-SA rats (Figure 7*B*). The GFP fluorescence levels in fixed sections visualized on a confocal microscope showed a high degree of overlap of cFos-GFP and endogenous immunolabeled Fos (Figure 7*D1,2*) consistent with other published work (Koya et al., 2012). A similar group of animals sacrificed immediately following the final extinction session (day 30) showed qualitatively similar GFP^+^ cells in living brain slices when evaluated on a confocal microscope (Figure 7*E1*). Cells from living brain slices collected at the same time point (day 30) were also suitable for high magnification *ex vivo* electrophysiology experiments (Figure 7*E2*).

Fluorescence-guided patch-clamp recordings of GFP^+/-^ cells were performed in slices from the cFos-GFP transgenic rats (Figure 7*F*) (Whitaker et al., 2017). Where possible, we attempted to sample equal numbers of GFP^+/-^ cells from each brain slice. In contrast to experiment 5, the final extinction or cue-test session was halted after 90 min to maximize the likelihood of a cFos-GFP signal. The firing responses of GFP^+^ and GFP ^―^ cells in the sal-Y and cocaine-SA groups after extinction differed (Figure 7*F*; two-way ANOVA with repeated measures of current, treatment *F*_(3,36)_ = 8.039, p = 0.0003; current: *F*_(7,252)_ = 8.971, p < 0.0001; interaction *F*_(21,252)_ = 3.188, p < 0.0001). A subsequent evaluation of only the sal-Y treatments showed that the number of action potentials increased as a function of increasing current injection, without detectable differences between GFP^+^ and GFP ^―^ cells (Figure 7*F*; two-way ANOVA with repeated measures of current GFP^+/-^ *F*_(1,18)_ = 2.731, p = 0.1157; current: *F*_(7,126)_ = 4.625, p < 0.0001; interaction *F*_(7,126)_ = 1.164, p = 0.3283). In contrast, a similar analysis of the cocaine-SA treatments showed GFP^+^ and GFP ^―^ cell responses to depolarizing current differed remarkably (Figure 7*F*; two-way ANOVA with repeated measures of current, GFP^+/-^ *F*_(1,18)_ = 15.51, p = 0.0010; current: *F*_(7,126)_ = 5.62, p < 0.0001; interaction *F*_(7,126)_ = 4.472, p < 0.0002). Even in brain slices from the same cocaine-SA with extinction animal, GFP^+^ cells from coca-SA with extinction rats showed on average a 3-4 fold enhancement of firing relative to the GFP ^―^ cells in the same animals (1 nA step, *t*_(144)_ = 4.996, p < 0.0001). The dot-plot summary for firing indicated that most cells (both GFP^+^ and GFP^-^) in sal-Y group displayed SFA after extinction (Figure 7*F*). By contrast, in the coca-SA group, most GFP^+^ cells lacked SFA, whereas the GFP^-^ cells in the same group were qualitatively similar to sal-Y responses after extinction (Figure 7*F*).

AHPs in GFP^+/-^ cells in the sal-Y and coca-SA treatment groups following extinction differed by treatment (two-way ANOVA, treatment *F*_(1,51)_ = 5.263, p = 0.0259; GFP^+/-^: *F*_(1,51)_ = 2.713, p = 0.1057; interaction *F*_(1,51)_ = 2.977, p = 0.0905). A follow up analysis showed that AHPs in GFP^+^ cells in the coca-SA group were reduced relative to the sal-Y group (Figure 7*F*; post hoc *t*_(51)_ = 2.904, p = 0.0108). In contrast, AHPs in GFP ― cells in the sal-Y and coca-SA group were similar (Figure 7*F*; post hoc *t*_(51)_ = 0.3940, p=0.9071). These data indicate that in animals with a history of cocaine-SA (but not sal-Y treatment), the expression of GFP^+^ overlaps with the persistent reduction in intrinsic inhibition after extinction training.

The effect of cocaine-predictive cues was similar to that observed above with extinction, however, GFP ^―^ cells were typically more difficult to find. After cue exposure, the firing responses of GFP^+/-^ cells in the sal-Y and cocaine-SA groups also differed (Figure 7*G*; two-way ANOVA with repeated measures of current, treatment *F*_(3,33)_ = 8.262, p = 0.0003; current: *F*_(7,231)_ = 46.18, p < 0.0001; interaction *F*_(21,231)_ = 9.144, p < 0.0001). In the sal-Y treatments, the number of action potentials increased as a function of increasing current injection, with no detectable difference between GFP^+^ and GFP ^―^ cells (Figure 7*G*; two-way ANOVA with repeated measures of current, GFP^+/-^ *F*_(1,16)_ = 0.4072, p = 0.5324; current: *F*_(7,112)_ = 4.311, p = 0.0003; interaction *F*_(7,112)_ = 0.4947, p = 0.8367). In contrast, in cocaine-SA treatments, the response of GFP^+^ and GFP ^―^ cell to depolarizing current differed and all GFP^+^ cells lacked SFA (Figure 7*F*; two-way ANOVA with repeated measures of current, GFP^+/-^ *F*_(1,17)_ = 8.947, p = 0.0082; current: *F*_(7,119)_ = 69.34, p < 0.0001; interaction *F*_(7,119)_ = 18.2, p < 0.0001). In the cocaine-SA group, GFP^+^ cells showed a 4 fold greater number of action potentials than in the GFP ^―^ cells (1 nA step, *t*_(136)_ = 7.414, p < 0.0001). After cue exposure, AHPs in GFP^+/-^ cells in the coca-SA and sal-Y treatments differed as a function of treatment (Figure 7*G*; two-way ANOVA, treatment *F*_(1,38)_ = 6.047, p = 0.0186; GFP^+/-^ *F*_(1,38)_ = 1.072, p < 0.3070; interaction *F*_(1,38)_ = 1.090, p < 0.3031). AHPs in GFP^+^ cells in the coca-SA group were reduced relative to the sal-Y group (post hoc *t*_(38)_ = 2.408, p = 0.0415), whereas AHPs in GFP ^―^ cells in the treatment groups were similar (Figure 7*G*; post hoc *t*_(38)_ = 1.031, p = 0.5227). Data from the extinction and cued-reinstatement experiments shown above indicate that the SFA ^―^cells from rats with a history of cocaine SA and conditioned cues also express cFos-GFP. In other studies, in behaviorally extinguished animals, GFP^+^ neurons comprising ensemble networks display strong activation in response to drug context environments (Koya et al., 2009; 2012). The nature of the Fos-generating stimulus in our behaviorally extinguished cocaine-SA animals remains to be determined.

Collectively, the above studies demonstrate that the reduction of intrinsic inhibition in a subpopulation of PL-PFC neurons after repeated operant coca-SA coincides temporally with changes in both drug seeking and neuronal activation. In summary, these experiments indicate that: 1) operant training for a drug (cocaine) or nondrug (sucrose) reinforcer is associated with neuromodification of two powerful types of intrinsic inhibition, and 2) whereas the modification for the nondrug reinforcer resolves during extinction, it persists for the drug reinforcer cocaine in a subpopulation of cells throughout extinction and cued-reinstatement of cocaine seeking.3) The persistent excitability is also associated with a Fos-GFP^+^ signature. Whether the activation of Fos reflects a re-exposure to the same environment where the animal first learned to associate the light and tone cues with cocaine or alternatively reflects competing activation of different ensembles (extinction versus cue) remains to be determined.

## Discussion

In rats, we found that learning to self-administer a drug reinforcer (cocaine) reduced PL-PFC intrinsic inhibition (as measured by SFA and AHPs) mediated by Kv7 channels. This modification persisted in a sub-population of PL-PFC neurons throughout extinction and was enhanced further by exposure to drug predictive cues. Such modifications were absent after similar treatments with a nondrug reinforcer (sucrose) or the same drug reinforcer (cocaine) administered passively under a yoked schedule. Cellular plasticity that is resistant to behavioral extinction and unusually robust is considered important for drug seeking behaviors. An existing prevailing theory is that drug-seeking stems from repeated intense neuronal activity that causes downstream biochemical changes in signal transduction, and thereby alters the responsiveness of cortical neurons to neurotransmitters that encode and interpret cues. In line with this, we found that dopamine regulation of intrinsic inhibition was occluded only in this hyper-responsive neural subpopulation and coincided with expression of the cFos-GFP transgene, suggesting prior robust dopamine transmission. Pharmacological stabilization of Kv7 channels with retigabine normalized SFA, potentiated *I*_Kv7_, and reduced reinstatement behavior. By contrast, cells lacking this modification showed normal levels of *I*_Kv7_ inhibition and insensitivity to retigabine, suggesting some pharmacological specificity for the activated subpopulation of cells mediating drug seeking. As Western blot analysis showed no overall changes in channel protein expression levels, it appears that repeated coca-SA causes a functional desensitization of inhibitory Kv7 channels. Given the role of SFA in synchronization of neuronal assemblies, this pathological shift away from intrinsic inhibition in this subpopulation of neurons may facilitate drug-seeking behavior.

### Role of intrinsic inhibition in cocaine self-administration and seeking

In primate PFC, learning to acquire a non-drug reward results in the formation of task-relevant neural ensembles (Buschman et al., 2012). The flexible nature of ensembles to form and reform underlies cognitive flexibility (Sejnowski and Paulsen, 2006; Womelsdorf and Fries, 2007). Chronic drug use impairs cognitive flexibility, rendering addicts unable to shift away from behavioral patterns learned during drug-taking (Briand et al., 2008; Kalivas and O’Brien, 2008; Porter et al., 2011; Schultz, 2011). SFA and AHPs are candidate cellular mechanisms that promote learning and memory by facilitating neuronal synchronization (Crook et al., 1998; Prescott and Sejnowski, 2008; Prescott et al., 2008; Sehgal et al., 2014). Transient reductions in SFA/AHP mechanisms by neuromodulators like dopamine accelerate the transition of LTD into LTP (Zaitsev and Anwyl, 2012). In other brain regions, reductions in intrinsic inhibition bias the recruitment of neuronal ensembles toward excitation and facilitate memory storage (Yiu et al., 2014) and possibly behavioral responses to drug associated cues.

Reductions in intrinsic inhibition are learning specific (Moyer et al., 1996; Thompson et al.,1996). Because such reductions are normally transient and normalize shortly after behavioral learning, the underlying mechanisms are considered essential for memory consolidation (Moyer et al., 1996). The re-stabilization or recovery of inhibition is equally important then in order to acquire new learning and extinguish a behavior (Kufahl et al., 2009; Pentkowski et al., 2012; 2014). Thus, persistent drug memories and craving could be associated with deficits in this process. In this regard, our finding that reinforcement learning for a drug reinforcer (cocaine) and a nondrug reinforcer (sucrose) differentially altered intrinsic inhibition is important. Drug and non-drug reinforcers differ in the degree to which they facilitate PFC dopamine release and form cue-reinforcement associations (Ciccocioppo et al., 2004; Tunstall and Kearns, 2014; Tunstall et al., 2014). Following repeated coca-SA, the combination of robust release of PFC dopamine and persistent unremitting reduction in intrinsic inhibition may bias cortical function toward future drug seeking.

### Select adaptations in a subpopulation of PL-PFC neurons

Previous studies indicate that sparse ensembles of neurons can encode a ‘seek’ rule learned by the repeated presentation of drug predictive cues and that selective ablation of these neurons reduces reinstatement for food and drug seeking (Bossert et al., 2012; Cruz et al., 2014), as well as incubation of drug craving (Fanous et al., 2012; Funk et al., 2016; Caprioli et al., 2017). These studies utilized cFos-GFP transgenic rat strategies, whereby the GFP signal serves as a proxy for the endogenous cFos gene product, and can also be targeted for selective inactivation by Duano2 (Cruz et al., 2013b). The Fos gene product is a known neuronal marker of intense firing activity (Morgan and Curran, 1991), and is activated by cocaine self-administration and context or cue-induced reinstatement (for review see (Cruz et al., 2015)). Fos expression requires robust elevations in calcium influx (McClung and Nestler, 2008) and associated with downstream biochemical changes that regulate gene expression (Cruz et al., 2013b).

Activity-induced increases in Fos are considered to represent the integration of neuronal activity driven by numerous synaptic alterations (Cifani et al., 2012; Koya et al., 2012; Whitaker et al., 2017; Ziminski et al., 2017a; 2017b). Using cFos-GFP transgenic rats, we observed that cocaine-SA training decreased intrinsic inhibition selectively in cFos/GFP^+^ neurons. Although the increased firing and increased Fos is logical, it contrasts with other reports showing reductions in excitatory synaptic strength (AMPAR/NMDAR current ratios) and glutamate release in cFos/GFP^+^ cells after stress-reinstatement of food seeking (Cifani et al., 2012; Koya et al., 2012). In those instances, the disparity in the cFos and excitation relationship was explained to reflect indirect homeostatic counteradaptations. Differences in those studies and our results may reflect differing reinforcers, brain regions, or the use of drug-associated cues. Irrespective, these studies and ours highlight the importance of studying functional changes, both synaptic and intrinsic, in behaviorally relevant neurons.

### The influence of dopamine

Dopamine exerts a multifaceted role in learning (Hollerman and Schultz, 1998), network connectivity (Eytan et al., 2004), and neuronal synchronization (Puig et al., 2014), including a stabilizing influence on cellular activity within neural ensembles (O’Donnell, 2003; Puig and Miller, 2012b; Miyawaki et al., 2014). Dopamine’s role in drug reward processing appears to facilitate the encoding of reward prediction, imprinting incentive value to reinforcers (energizing approach behavior) and facilitating learning of reward associations (conditioning) through its modulation of subcortical (including the nucleus accumbens) and cortical brain regions (Di Chiara, 1999; Schultz, 1999; Waelti et al., 2001; Schultz, 2002; Redish, 2004; Steinberg et al., 2013). Cellular adaptations in dopaminergic signaling contribute to relapse to drug seeking (Hyman et al., 2006), in part by disrupting transmission in the prefrontal cortex (Anderson and Pierce, 2005; Dong et al., 2005a; Huang et al., 2007; Sidiropoulou et al., 2009; Buchta and Riegel, 2015). In tissue from rats with a history of repeated coca-SA training, we observed a decreased ability of dopamine to alter firing and *I*_Kv7_ in a subpopulation of cells, which may reflect prior sensitization or overactivation in this neuronal population. Unexpectedly, the SFA^+^ population of cells from both the sal-Y and the coca-SA treatments showed normal responses to dopamine, indicating some neuronal selectivity in the modification of intrinsic inhibition. The reason for this selectivity is unclear but may reflect differences in dopamine receptor expression. Chronic exposure to cocaine causes rapid, robust and persistent increases in D1 receptor-mediated signaling that result in a prominent upregulation of Fos (Henry and White, 1995; McClung and Nestler, 2003). Functional D1 receptors are required for Fos induction by cocaine (Drago et al., 1996; Moratalla et al., 1996; Zhang et al., 2002; 2004), and cocaine-induced behaviors are greatly reduced in transgenic mice with inactivated Fos in D1-receptor expressing cells throughout the brain (Zhang et al., 2006). In SFA^-^ cells, we observed both occluded dopamine responses and a strong association with the cFos-GFP transgene (even under extinction-like conditions). Perhaps prior robust overactivation or sensitization of dopamine signaling functionally reduces intrinsic inhibition and further modulation by dopamine in these neurons.

### Kv7 ion channels

Why Kv7 channel currents are reduced following cocaine-SA is unclear. Cocaine exposure is known to elevate levels of intracellular calcium, which reportedly can initiate a well-described calmodulin-sensitive suppression of the Kv7 ion channel activity, resulting in hyper-excitability (Kirkwood et al., 1991; Selyanko and Brown, 1996; Cruzblanca et al., 1998; Gamper and Shapiro, 2003; Gamper et al., 2003). Other possible mechanisms regulating the channel will also need to be examined including elevated cAMP and PKA activity (Dong et al., 2005b; Nasif et al., 2005; Ford et al., 2009), and/or reduced levels of PtdIns(4,5)P2 (Park et al., 2001; Zhang and Linden, 2003; Li et al., 2005). The adaptations outlined above all likely contribute to increased cue-induced drug-seeking behavior long after cessation of drug use. Nevertheless, our results demonstrate a functional role for Kv7 channel mediated inhibition in a behaviorally relevant subpopulation of PL-PFC neurons important for cue-induced reinstatement of drug seeking. Future studies combining pharmacological treatments like retigabine with newer strategies for time-sensitive labeling of behaviorally relevant neurons using cFos-TetTag in transgenic mice (Reijmers et al., 2007) may further refine precisely when these adaptations disrupt Kv7 channel function and how this contributes to cue-induced cocaine seeking.

## Acknowledgements

NIDA grants F31-DA036989 (W.C.B.), T32-DA007288 (J.P.C; W.C.B.; PD: J.F.M.), R01-DA027664 (PI: C.W),R01-NS098772 (PI: R.K.), R01-DA042852 (PI: R.K.), P50-DA015369 (PI: A.C.R.; PD: P.W.K.), and R01-DA033342A (A.C.R.)

## Notes

Conflict of interest: The authors declare no competing financial interests.

## References

Aiken SP, Lampe BJ, Murphy PA, Brown BS (1995) Reduction of spike frequency adaptation and blockade of M-current in rat CA1 pyramidal neurones by linopirdine (DuP 996), a neurotransmitter release enhancer. Br J Pharmacol 115:1163–1168.

Anderson SM, Pierce RC (2005) Cocaine-induced alterations in dopamine receptor signaling: implications for reinforcement and reinstatement. Pharmacol Ther 106:389–403.

Battefeld A, Tran BT, Gavrilis J, Cooper EC, Kole MHP (2014) Heteromeric Kv7.2/7.3 Channels Differentially Regulate Action Potential Initiation and Conduction in Neocortical Myelinated Axons. J Neurosci 34:3719–3732.

Benda J, Longtin A, Maler L (2005) Spike-frequency adaptation separates transient communication signals from background oscillations. J Neurosci 25:2312–2321.

Bobadilla JA, Heinsbroek A-C, Gipson CD, Griffin WC, Fowler CD, Kenny PJ, Kalivas PW (2017) Corticostriatal plasticity, neuronal ensembles, and regulation of drug-seeking behavior. Prog Brain Res 235:93–112.

Bossert JM, Marchant NJ, Calu DJ, Shaham Y (2013) The reinstatement model of drug relapse: recent neurobiological findings, emerging research topics, and translational research. Psychopharmacology 229:453–476.

Bossert JM, Stern AL, Theberge FRM, Cifani C, Koya E, Hope BT, Shaham Y (2011) Ventral medial prefrontal cortex neuronal ensembles mediate context-induced relapse to heroin. Nat Neurosci 14:420–422.

Bossert JM, Stern AL, Theberge FRM, Marchant NJ, Wang H-L, Morales M, Shaham Y (2012) Role of projections from ventral medial prefrontal cortex to nucleus accumbens shell in context-induced reinstatement of heroin seeking. J Neurosci 32:4982–4991.

Briand LA, Flagel SB, Garcia-Fuster MJ, Watson SJ, Akil H, Sarter M, Robinson TE (2008) Persistent alterations in cognitive function and prefrontal dopamine D2 receptors following extended, but not limited, access to self-administered cocaine. Neuropsychopharmacology 33:2969–2980.

Brown DA, Passmore GM (2009) Neural KCNQ (Kv7) channels. Br J Pharmacol 156:1185–1195.

Buchta WC, Mahler SV, Harlan B, Aston-Jones GS, Riegel AC (2017) Dopamine terminals from the ventral tegmental area gate intrinsic inhibition in the prefrontal cortex. Physiological Reports 5:e13198.

Buchta WC, Riegel AC (2015) Chronic cocaine disrupts mesocortical learning mechanisms. Brain Res 1628:88–103.

Buschman TJ, Denovellis EL, Diogo C, Bullock D, Miller EK (2012) Synchronous Oscillatory Neural Ensembles for Rules in the Prefrontal Cortex. Neuron 76:838–846.

Caprioli D, Venniro M, Zhang M, Bossert JM, Warren BL, Hope BT, Shaham Y (2017) Role of Dorsomedial Striatum Neuronal Ensembles in Incubation of Methamphetamine Craving after Voluntary Abstinence. J Neurosci 37:1014–1027.

Ceci A, Brambilla A, Duranti P, Grauert M, Grippa N, Borsini F (1999) Effect of antipsychotic drugs and selective dopaminergic antagonists on dopamine-induced facilitatory activity in prelimbic cortical pyramidal neurons. An in vitro study. Neuroscience 93:107–115.

Ciccocioppo R, Martin-Fardon R, Weiss F (2004) Stimuli associated with a single cocaine experience elicit long-lasting cocaine-seeking. Nat Neurosci 7:495–496.

Cifani C, Koya E, Navarre BM, Calu DJ, Baumann MH, Marchant NJ, Liu Q-R, Khuc T, Pickel J, Lupica CR, Shaham Y, Hope BT (2012) Medial prefrontal cortex neuronal activation and synaptic alterations after stress-induced reinstatement of palatable food seeking: a study using c-fos-GFP transgenic female rats. J Neurosci 32:8480–8490.

Coulter DA, Turco Lo JJ, Kubota M, Disterhoft JF, Moore JW, Alkon DL (1989) Classical conditioning reduces amplitude and duration of calcium-dependent afterhyperpolarization in rabbit hippocampal pyramidal cells. J Neurophysiol 61:971–981.

Crook SM, Ermentrout GB, Bower JM (1998) Spike frequency adaptation affects the synchronization properties of networks of cortical oscillations. Neural Comput 10:837–854.

Cruz FC, Babin KR, Leao RM, Goldart EM, Bossert JM, Shaham Y, Hope BT (2014) Role of Nucleus Accumbens Shell Neuronal Ensembles in Context-Induced Reinstatement of Cocaine-Seeking. J Neurosci 34:7437–7446.

Cruz FC, Koya E, Guez-Barber DH, Bossert JM, Lupica CR, Shaham Y, Hope BT (2013a) New technologies for examining the role of neuronal ensembles in drug addiction and fear. Nat Rev Neurosci 14:743–754.

Cruz FC, Koya E, Guez-Barber DH, Bossert JM, Lupica CR, Shaham Y, Hope BT (2013b) New technologies for examining the role of neuronal ensembles in drug addiction and fear. Nat Rev Neurosci 14:743–754 Available at: http://dx.doi.org/10.1038/nrn3597.

Cruz FC, Rubio FJ, Hope BT (2015) Using c-fos to study neuronal ensembles in corticostriatal circuitry of addiction. Brain Res 1628:157–173.

Cruzblanca H, Koh DS, Hille B (1998) Bradykinin inhibits M current via phospholipase C and Ca2+ release from IP3-sensitive Ca2+ stores in rat sympathetic neurons. Proc Natl Acad Sci USA 95:7151–7156.

Delmas P, Brown DA (2005) Pathways modulating neural KCNQ/M (Kv7) potassium channels. Nat Rev Neurosci 6:850–862.

Di Chiara G (1999) Drug addiction as dopamine-dependent associative learning disorder. Eur J Pharmacol 375:13–30.

Dong Y, Nasif FJ, Tsui JJ, Ju WY, Cooper DC, Hu X-T, Malenka RC, White FJ (2005a) Cocaine-induced plasticity of intrinsic membrane properties in prefrontal cortex pyramidal neurons: adaptations in potassium currents. J Neurosci 25:936–940.

Dong Y, Nasif FJ, Tsui JJ, Ju WY, Cooper DC, Hu X-T, Malenka RC, White FJ (2005b) Cocaine-induced plasticity of intrinsic membrane properties in prefrontal cortex pyramidal neurons: adaptations in potassium currents. J Neurosci 25:936–940.

Drago J, Gerfen CR, Westphal H, Steiner H (1996) D1 dopamine receptor-deficient mouse: cocaine-induced regulation of immediate-early gene and substance P expression in the striatum. Neuroscience 74:813–823.

Epstein DH, Preston KL, Stewart J, Shaham Y (2006) Toward a model of drug relapse: an assessment of the validity of the reinstatement procedure. Psychopharmacology 189:1–16.

Euston DR, Gruber AJ, McNaughton BL (2012) The Role of Medial Prefrontal Cortex in Memory and Decision Making. Neuron 76:1057–1070.

Eytan D, Minerbi A, Ziv N, Marom S (2004) Dopamine-induced dispersion of correlations between action potentials in networks of cortical neurons. J Neurophysiol 92:1817–1824.

Faber ES, Callister RJ, Sah P (2001) Morphological and electrophysiological properties of principal neurons in the rat lateral amygdala in vitro. J Neurophysiol 85:714–723.

Fanous S, Goldart EM, Theberge FRM, Bossert JM, Shaham Y, Hope BT (2012) Role of orbitofrontal cortex neuronal ensembles in the expression of incubation of heroin craving. J Neurosci 32:11600–11609.

Ford KA, Wolf ME, Hu X-T (2009) Plasticity of L-type Ca2+ channels after cocaine withdrawal. Synapse 63:690–697.

Funk D, Coen K, Tamadon S, Hope BT, Shaham Y, Lê AD (2016) Role of Central Amygdala Neuronal Ensembles in Incubation of Nicotine Craving. J Neurosci 36:8612–8623.

Gamper N, Shapiro MS (2003) Calmodulin mediates Ca2+-dependent modulation of M-type K+ channels. J Gen Physiol 122:17–31.

Gamper N, Stockand JD, Shapiro MS (2003) Subunit-specific modulation of KCNQ potassium channels by Src tyrosine kinase. J Neurosci 23:84–95.

Gipson CD, Kupchik YM, Kalivas PW (2014) Rapid, transient synaptic plasticity in addiction. Neuropharmacology 76 Pt B:276–286.

Gipson CD, Kupchik YM, Shen H, Reissner KJ, Thomas CA, Kalivas PW (2013) Relapse induced by cues predicting cocaine depends on rapid, transient synaptic potentiation. Neuron 77:867–872.

Gorelova N, Seamans JK, Yang CR (2002) Mechanisms of dopamine activation of fast-spiking interneurons that exert inhibition in rat prefrontal cortex. J Neurophysiol 88:3150–3166.

Graybiel AM (2008) Habits, rituals, and the evaluative brain. Annu Rev Neurosci 31:359–387.

Gu N, Vervaeke K, Hu H, Storm JF (2005) Kv7/KCNQ/M and HCN/h, but not KCa2/SK channels, contribute to the somatic medium after-hyperpolarization and excitability control in CA1 hippocampal pyramidal cells. J Physiol (Lond) 566:689–715.

Gulledge AT, Jaffe DB (1998) Dopamine Decreases the Excitability of Layer V Pyramidal Cells in the Rat Prefrontal Cortex. Journal of Neuroscience 18:9139–9151.

Henry DJ, White FJ (1995) The persistence of behavioral sensitization to cocaine parallels enhanced inhibition of nucleus accumbens neurons. Journal of Neuroscience 15:6287–6299.

Henze DA, González-Burgos GR, Urban NN, Lewis DA, Barrionuevo G (2000) Dopamine increases excitability of pyramidal neurons in primate prefrontal cortex. J Neurophysiol 84:2799–2809.

Herrera D, Robertson H (1996) Activation of in the brain. Prog Neurobiol 50:83–107.

Hollerman JR, Schultz W (1998) Dopamine neurons report an error in the temporal prediction of reward during learning. Nat Neurosci 1:304–309.

Huang C-C, Yang P-C, Lin H-J, Hsu K-S (2007) Repeated cocaine administration impairs group II metabotropic glutamate receptor-mediated long-term depression in rat medial prefrontal cortex. J Neurosci 27:2958–2968.

Huang H, Trussell LO (2011) KCNQ5 channels control resting properties and release probability of a synapse. Nat Neurosci 14:840–847.

Hyman SE, Malenka RC, Nestler EJ (2006) Neural mechanisms of addiction: the role of reward-related learning and memory. Annu Rev Neurosci 29:565–598.

Johnston CA, Watts VJ (2003) Sensitization of adenylate cyclase: a general mechanism of neuroadaptation to persistent activation of Galpha(i/o)-coupled receptors? Life Sci 73:2913–2925.

Kalivas PW, McFarland K (2003) Brain circuitry and the reinstatement of cocaine-seeking behavior. Psychopharmacology 168:44–56.

Kalivas PW, O’Brien C (2008) Drug addiction as a pathology of staged neuroplasticity. Neuropsychopharmacology 33:166–180.

Kirkwood A, Simmons MA, Mather RJ, Lisman J (1991) Muscarinic suppression of the M-current is mediated by a rise in internal Ca2+ concentration. Neuron 6:1009–1014.

Koya E, Cruz FC, Ator R, Golden SA, Hoffman AF, Lupica CR, Hope BT (2012) Silent synapses in selectively activated nucleus accumbens neurons following cocaine sensitization. Nat Neurosci 15:1556–1562.

Koya E, Golden SA, Harvey BK, Guez-Barber DH, Berkow A, Simmons DE, Bossert JM, Nair SG, Uejima JL, Marin MT, Mitchell TB, Farquhar D, Ghosh SC, Mattson BJ, Hope BT (2009) Targeted disruption of cocaine-activated nucleus accumbens neurons prevents context-specific sensitization. Nat Neurosci 12:1069–1073.

Kufahl PR, Zavala AR, Singh A, Thiel KJ, Dickey ED, Joyce JN, Neisewander JL (2009) c-Fos expression associated with reinstatement of cocaine-seeking behavior by response-contingent conditioned cues. Synapse 63:823–835.

Lancaster B, Nicoll RA (1987) Properties of two calcium-activated hyperpolarizations in rat hippocampal neurones. J Physiol (Lond) 389:187–203.

Lawrence JJ, Saraga F, Churchill JF, Statland JM, Travis KE, Skinner FK, McBain CJ (2006) Somatodendritic Kv7/KCNQ/M channels control interspike interval in hippocampal interneurons. J Neurosci 26:12325–12338.

Lerman DC, Iwata BA (1995) Prevalence of the extinction burst and its attenuation during treatment. J Appl Behav Anal 28:93–94.

Li Y, Gamper N, Hilgemann DW, Shapiro MS (2005) Regulation of Kv7 (KCNQ) K+ channel open probability by phosphatidylinositol 4,5-bisphosphate. J Neurosci 25:9825–9835.

Madison DV, Nicoll RA (1982) Noradrenaline blocks accommodation of pyramidal cell discharge in the hippocampus. Nature 299:636–638.

Madison DV, Nicoll RA (1984) Control of the repetitive discharge of rat CA 1 pyramidal neurones in vitro. J Physiol (Lond) 354:319–331.

Malenka RC, Nicoll RA (1986) Dopamine decreases the calcium-activated afterhyperpolarization in hippocampal CA1 pyramidal cells. Brain Res 379:210–215.

Matsumoto M, Hikosaka O (2009) Two types of dopamine neuron distinctly convey positive and negative motivational signals. Nature 459:837–841.

McClung CA, Nestler EJ (2003) Regulation of gene expression and cocaine reward by CREB and DeltaFosB. Nat Neurosci 6:1208–1215.

McClung CA, Nestler EJ (2008) Neuroplasticity mediated by altered gene expression. Neuropsychopharmacology 33:3–17.

McGlinchey EM, James MH, Mahler SV, Pantazis C, Aston-Jones G (2016) Prelimbic to Accumbens Core Pathway Is Recruited in a Dopamine-Dependent Manner to Drive Cued Reinstatement of Cocaine Seeking. J Neurosci 36:8700–8711.

McKay BM, Matthews EA, Oliveira FA, Disterhoft JF (2009) Intrinsic neuronal excitability is reversibly altered by a single experience in fear conditioning. J Neurophysiol 102:2763–2770.

McLaughlin J, See RE (2003) Selective inactivation of the dorsomedial prefrontal cortex and the basolateral amygdala attenuates conditioned-cued reinstatement of extinguished cocaine-seeking behavior in rats. Psychopharmacology 168:57–65.

Miyawaki T, Norimoto H, Ishikawa T, Watanabe Y, Matsuki N, Ikegaya Y (2014) Dopamine receptor activation reorganizes neuronal ensembles during hippocampal sharp waves in vitro. PLoS ONE 9:e104438.

Moratalla R, Xu M, Tonegawa S, Graybiel AM (1996) Cellular responses to psychomotor stimulant and neuroleptic drugs are abnormal in mice lacking the D1 dopamine receptor. Proc Natl Acad Sci USA 93:14928–14933.

Morgan JI, Cohen DR, Hempstead JL, Curran T (1987) Mapping patterns of c-fos expression in the central nervous system after seizure. Science 237:192–197.

Morgan JI, Curran T (1988) Calcium as a modulator of the immediate-early gene cascade in neurons. Cell Calcium 9:303–311.

Morgan JI, Curran T (1991) Proto-oncogene transcription factors and epilepsy. Trends Pharmacol Sci 12:343–349.

Moyer JR, Power JM, Thompson LT, Disterhoft JF (2000) Increased excitability of aged rabbit CA1 neurons after trace eyeblink conditioning. Journal of Neuroscience 20:5476–5482.

Moyer JR, Thompson LT, Disterhoft JF (1996) Trace eyeblink conditioning increases CA1 excitability in a transient and learning-specific manner. Journal of Neuroscience 16:5536–5546.

Nasif FJ, FJ Hu X-T, White (2005) Repeated cocaine administration increases voltage-sensitive calcium currents in response to membrane depolarization in medial prefrontal cortex pyramidal neurons. J Neurosci 25:3674–3679.

O’Donnell P (2003) Dopamine gating of forebrain neural ensembles. Eur J Neurosci 17:429–435.

Oh MM, Disterhoft JF (2015) Increased Excitability of Both Principal Neurons and Interneurons during Associative Learning. The Neuroscientist 21:372–384.

Park SJ, Itoh T, Takenawa T (2001) Phosphatidylinositol 4-phosphate 5-kinase type I is regulated through phosphorylation response by extracellular stimuli. J Biol Chem 276:4781–4787.

Pentkowski NS, Cheung THC, Toy WA, Adams MD, Neumaier JF, Neisewander JL (2012) Protracted Withdrawal from Cocaine Self-Administration Flips the Switch on 5-HT1B Receptor Modulation of Cocaine Abuse-Related Behaviors. Biol Psychiatry 72:396–404.

Pentkowski NS, Harder BG, Brunwasser SJ, Bastle RM, Peartree NA, Yanamandra K, Adams MD, T Der-Ghazarian, Neisewander JL (2014) Pharmacological evidence for an abstinence-induced switch in 5-HT1B receptor modulation of cocaine self-administration and cocaine-seeking behavior. ACS Chem Neurosci 5:168–176.

Pinto L, Dan Y (2015) Cell-Type-Specific Activity in Prefrontal Cortex during Goal-Directed Behavior. Neuron 87:437–450.

Porter JN, Olsen AS, Gurnsey K, Dugan BP, Jedema HP, Bradberry CW (2011) Chronic cocaine self-administration in rhesus monkeys: impact on associative learning, cognitive control, and working memory. J Neurosci 31:4926–4934.

Prescott SA, Ratté S, de Koninck Y, Sejnowski TJ (2008) Pyramidal neurons switch from integrators in vitro to resonators under in vivo-like conditions. J Neurophysiol 100:3030–3042.

Prescott SA, Sejnowski TJ (2008) Spike-rate coding and spike-time coding are affected oppositely by different adaptation mechanisms. J Neurosci 28:13649–13661.

Puig MV, Antzoulatos EG, Miller EK (2014) Prefrontal dopamine in associative learning and memory. Neuroscience 282C:217–229.

Puig MV, Miller EK (2012a) The role of prefrontal dopamine D1 receptors in the neural mechanisms of associative learning. Neuron 74:874–886.

Puig MV, Miller EK (2012b) The Role of Prefrontal Dopamine D1 Receptors in the Neural Mechanisms of Associative Learning. Neuron 74:874–886.

Redish AD (2004) Addiction as a computational process gone awry. Science 306:1944–1947.

Reijmers LG, Perkins BL, Matsuo N, Mayford M (2007) Localization of a stable neural correlate of associative memory. Science 317:1230–1233.

Riegel AC, Williams JT (2008) CRF facilitates calcium release from intracellular stores in midbrain dopamine neurons. Neuron 57:559–570.

Rosenkranz JA, Grace AA (2002) Cellular mechanisms of infralimbic and prelimbic prefrontal cortical inhibition and dopaminergic modulation of basolateral amygdala neurons in vivo. J Neurosci 22:324–337.

Schultz W (1999) The Reward Signal of Midbrain Dopamine Neurons. News Physiol Sci 14:249–255.

Schultz W (2011) Potential vulnerabilities of neuronal reward, risk, and decision mechanisms to addictive drugs. Neuron 69:603–617.

See RE (2009) Dopamine D1 receptor antagonism in the prelimbic cortex blocks the reinstatement of heroin-seeking in an animal model of relapse. Int J Neuropsychopharmacol 12:431–436.

Sehgal M, Ehlers VL, Moyer JR (2014) Learning enhances intrinsic excitability in a subset of lateral amygdala neurons. Learn Mem 21:161–170.

Sejnowski TJ, Paulsen O (2006) Network oscillations: emerging computational principles. J Neurosci 26:1673–1676.

Selyanko AA, Brown DA (1996) Intracellular calcium directly inhibits potassium M channels in excised membrane patches from rat sympathetic neurons. Neuron 16:151–162.

Shah MM, Javadzadeh-Tabatabaie M, Benton DCH, Ganellin CR, Haylett DG (2006) Enhancement of hippocampal pyramidal cell excitability by the novel selective slow-afterhyperpolarization channel blocker 3-(triphenylmethylaminomethyl)pyridine (UCL2077). Mol Pharmacol 70:1494–1502.

Shaham Y, Shalev U, Lu L, de Wit H, Stewart J (2003) The reinstatement model of drug relapse: history, methodology and major findings. Psychopharmacology 168:3–20.

Sidiropoulou K, Lu F-M, Fowler MA, Xiao R, Phillips C, Ozkan ED, Zhu MX, White FJ, Cooper DC (2009) Dopamine modulates an mGluR5-mediated depolarization underlying prefrontal persistent activity. Nat Neurosci 12:190–199.

Simon Peter Peron FG (2009) Role of spike-frequency adaptation in shaping neuronal response to dynamic stimuli. Biological cybernetics 100:505–520.

Smith ACW, Kupchik YM, Scofield MD, Gipson CD, Wiggins A, Thomas CA, Kalivas PW (2014) Synaptic plasticity mediating cocaine relapse requires matrix metalloproteinases. Nat Neurosci 17:1655–1657.

Song C, Ehlers VL, Moyer JR (2015) Trace Fear Conditioning Differentially Modulates Intrinsic Excitability of Medial Prefrontal Cortex-Basolateral Complex of Amygdala Projection Neurons in Infralimbic and Prelimbic Cortices. J Neurosci 35:13511–13524.

Spencer S, Garcia-Keller C, Roberts-Wolfe D, Heinsbroek JA, Mulvaney M, Sorrell A, Kalivas PW (2017) Cocaine Use Reverses Striatal Plasticity Produced During Cocaine Seeking. Biol Psychiatry 81:616–624.

Stefanik MT, Kupchik YM, Kalivas PW (2016) Optogenetic inhibition of cortical afferents in the nucleus accumbens simultaneously prevents cue-induced transient synaptic potentiation and cocaine-seeking behavior. Brain structure & function 221:1681–1689.

Steinberg EE, Keiflin R, Boivin JR, Witten IB, Deisseroth K, Janak PH (2013) A causal link between prediction errors, dopamine neurons and learning. Nat Neurosci 16:966–973.

Steketee JD, Kalivas PW (2011) Drug wanting: behavioral sensitization and relapse to drug-seeking behavior. Pharmacol Rev 63:348–365.

Storm JF (1990) Potassium currents in hippocampal pyramidal cells. Prog Brain Res 83:161–187.

Taniguchi M, Carreira MB, Cooper YA, Bobadilla A-C, Heinsbroek JA, Koike N, Larson EB, Balmuth EA, Hughes BW, Penrod RD, Kumar J, Smith LN, Guzman D, Takahashi JS, Kim T-K, Kalivas PW, Self DW, Lin Y, Cowan CW (2017) HDAC5 and Its Target Gene, Npas4, Function in the Nucleus Accumbens to Regulate Cocaine-Conditioned Behaviors. Neuron 96:130–144.e136.

Thompson LT, Moyer JR, Disterhoft JF (1996) Transient changes in excitability of rabbit CA3 neurons with a time course appropriate to support memory consolidation. J Neurophysiol 76:1836–1849.

Tunstall BJ, Kearns DN (2014) Reinstatement in a cocaine versus food choice situation: reversal of preference between drug and non-drug rewards. Addict Biol 19:838–848.

Tunstall BJ, Kearns DN (2016) Cocaine can generate a stronger conditioned reinforcer than food despite being a weaker primary reinforcer. Addict Biol 21:282–293.

Tunstall BJ, Riley AL, Kearns DN (2014) Drug specificity in drug versus food choice in male rats. Experimental and clinical psychopharmacology 22:364–372.

Venniro M, Caprioli D, Shaham Y (2016) Animal models of drug relapse and craving: From drug priming-induced reinstatement to incubation of craving after voluntary abstinence. Prog Brain Res 224:25–52 Available at: http://eutils.ncbi.nlm.nih.gov/entrez/eutils/elink.fcgi?dbfrom=pubmed&id=26822352&retmode=ref&cmd=prli nks.

Waelti P, Dickinson A, Schultz W (2001) Dopamine responses comply with basic assumptions of formal learning theory. Nature 412:43–48.

Wang HS, Pan Z, Shi W, Brown BS, Wymore RS, Cohen IS, Dixon JE, McKinnon D (1998) KCNQ2 and KCNQ3 potassium channel subunits: molecular correlates of the M-channel. Science 282:1890–1893.

Whitaker LR, Warren BL, Venniro M, Harte TC, McPherson KB, Beidel J, Bossert JM, Shaham Y, Bonci A, Hope BT (2017) Bidirectional Modulation of Intrinsic Excitability in Rat Prelimbic Cortex Neuronal Ensembles and Non-Ensembles after Operant Learning. J Neurosci 37:8845–8856.

Williams CL, Buchta WC, Riegel AC (2014) CRF-R2 and the heterosynaptic regulation of VTA glutamate during reinstatement of cocaine seeking. J Neurosci 34:10402–10414 Available at: http://www.jneurosci.org/cgi/doi/10.1523/JNEUROSCI.0911-13.2014.

Williams MJ, Adinoff B (2007) The Role of Acetylcholine in Cocaine Addiction. Neuropsychopharmacology 33:1779–1797.

Womelsdorf T, Fries P (2007) The role of neuronal synchronization in selective attention. Curr Opin Neurobiol 17:154–160.

Yang CR, Seamans JK (1996) Dopamine D1 receptor actions in layers V-VI rat prefrontal cortex neurons in vitro: modulation of dendritic-somatic signal integration. Journal of Neuroscience 16:1922–1935.

Yiu AP, Mercaldo V, Yan C, Richards B, Rashid AJ, Hsiang H-LL, Pressey J, Mahadevan V, Tran MM, Kushner SA, Woodin MA, Frankland PW, Josselyn SA (2014) Neurons Are Recruited to a Memory Trace Based on Relative Neuronal Excitability Immediately before Training. Neuron 83:722–735.

Zaitsev AV, Anwyl R (2012) Inhibition of the slow afterhyperpolarization restores the classical spike timing-dependent plasticity rule obeyed in layer 2/3 pyramidal cells of the prefrontal cortex. J Neurophysiol 107:205–215.

Zhang D, Zhang L, Lou DW, Nakabeppu Y, Zhang J, Xu M (2002) The dopamine D1 receptor is a critical mediator for cocaine-induced gene expression. J Neurochem 82:1453–1464.

Zhang J, Zhang L, Jiao H, Zhang Q, Zhang D, Lou D, Katz JL, Xu M (2006) c-Fos facilitates the acquisition and extinction of cocaine-induced persistent changes. J Neurosci 26:13287–13296.

Zhang L, Lou D, Jiao H, Zhang D, Wang X, Xia Y, Zhang J, Xu M (2004) Cocaine-induced intracellular signaling and gene expression are oppositely regulated by the dopamine D1 and D3 receptors. J Neurosci 24:3344–3354.

Zhang W, Linden DJ (2003) The other side of the engram: experience-driven changes in neuronal intrinsic excitability. Nat Rev Neurosci 4:885–900.

Zheng P, Zhang XX, Bunney BS, Shi WX (1999) Opposite modulation of cortical N-methyl-D-aspartate receptor-mediated responses by low and high concentrations of dopamine. Neuroscience 91:527–535.

Ziminski JJ, Hessler S, Margetts-Smith G, Sieburg MC, Crombag HS, Koya E (2017a) Changes in Appetitive Associative Strength Modulates Nucleus Accumbens, But Not Orbitofrontal Cortex Neuronal Ensemble Excitability. J Neurosci 37:3160–3170.

Ziminski JJ, Sieburg MC, Margetts-Smith G, Crombag HS, Koya E (2017b) Regional Differences in Striatal Neuronal Ensemble Excitability Following Cocaine and Extinction Memory Retrieval in Fos-GFP Mice. Neuropsychopharmacology 18:10579.

